# Essential and dual effects of Notch activity on a natural transdifferentiation event

**DOI:** 10.1101/2024.09.11.612396

**Authors:** Thomas Daniele, Jeanne Cury, Marie-Charlotte Morin, Arnaud Ahier, Davide Isaia, Sophie Jarriault

## Abstract

Cell identity can be reprogrammed, naturally or experimentally, albeit with low frequency. Why given cells, but not their neighbours, undergo a cell identity conversion remains unclear. We find that Notch signalling plays a key role to promote natural transdifferentiation in *C. elegans*. Endogenous Notch signal endows a cell with the competence to transdifferentiate by promoting plasticity factors expression (*hlh-16/Olig* and *sem-4/Sall*). Strikingly, exogenous Notch can trigger ectopic transdifferentiation *in vivo*. However, Notch signalling can both promote and block transdifferentiation depending on its activation timing. Notch only promotes transdifferentiation during an early precise window of opportunity and signal duration must be tightly controlled in time. Our findings emphasise the importance of temporality and dynamics of the underlying molecular events preceding the initiation of natural cell reprogramming. Finally, our results support a model where both an extrinsic signal and the intrinsic cellular context combine to empower a cell with the competence to transdifferentiate.

## Introduction

During metazoan development, cells become increasingly specialised as they acquire a final identity that fulfils a specific function in a particular tissue or organ. Nevertheless differentiated cells have been shown to naturally change their identity in many different species and tissues^1,2^, although demonstration at the single cell level is often lacking. The process whereby a cell stably and fully switches from one differentiated identity to another has been called transdifferentiation (Td) or direct reprogramming^3,4^. Examples include the conversion of pigmented cells into lens cells during lens regeneration in different species^5^, the conversion of kidney distal tubule epithelial cells into an endocrine gland in zebrafish^6^, or, in mammals, the conversion of heart venous cells into coronary artery cells^7^.

Direct reprogramming can be experimentally reproduced by forcing the expression of specific transcription factors in differentiated cells *in vivo* or *in vitro*^8–10^. Experimental models of induced Td typically exhibit a fairly low success rate. Thus, within a population of similar cells, only a few will actually switch identity. Most induced cells will not undergo Td, making it difficult to focus on the mechanisms involved, let alone predict which cells will Td. Furthermore, a number of studies have shown that for a given inducing cue, certain cells types can be reprogrammed but not others^11–13^. These pioneer studies on natural and induced direct reprogramming have highlighted important questions that remain unanswered. Why is only a small subset of cells within a population able to transdifferentiate? How is a cell endowed with the capability of changing its identity, and particularly for natural Td events, to what extent do extrinsic cues and context, such as the micro-environment, or the intrinsic cellular content, modulate this capability? Here we have taken advantage of a single-celled Td event, which naturally occurs in 100% of animals and can be followed *in vivo*, to address these issues. We report evidence that an extrinsic signal through the Notch pathway is involved in potentiating *in vivo* Td by promoting intrinsic reprogramming factor expression.

The Notch signalling pathway is a highly conserved pathway involved in a plethora of essential developmental processes including cellular plasticity. It plays a role in cell fate determination and differentiation or tissue homeostasis by regulating apoptosis or stem cell maintenance^14,15^ and mediates both lateral or inductive signalling^16^. Notch signalling studies have been pioneered in invertebrate model systems such as *Drosophila melanogaster* and the nematode worm *Caenorhabditis elegans* where the Notch receptor was first identified^17–20^. Mechanistic aspects of Notch signal transduction, which involves the interaction between a receptor-expressing cell and a ligand-expressing cell, are well understood^21^. The contact between the Notch receptor(s) and its ligand(s) induces a succession of proteolytic cleavages of the receptor that lead to the release of the Notch intracellular domain (NICD). The NICD translocates to the nucleus and then interacts with the DNA-binding protein CSL (CBF1/RBPjκ/Su(H)/Lag-1) to activate the transcription of target genes^21–27^.

Notch signalling has been shown to be necessary and sufficient to make a rectal cell named “Y”, in *C. elegans*^17^. In absence of Notch signal, no Y cell is made^17,28^. The Y cell is one of six fully differentiated rectal cells forming the rectal tube, and the only one with an extraordinary behaviour: it subsequently migrates away from the rectum during mid-larval development and switches its identity to become a motoneuron, named PDA^28,29^ (Fig1A). This process has been the focus of in-depth studies at the single cell level and unambiguously shown to be a stereotyped *bona fide* Td event^28,30–33^. Intriguingly, the other rectal neighbouring cells, seemingly identical, never change their identity: what makes the Y cell special?

**Figure 1:**
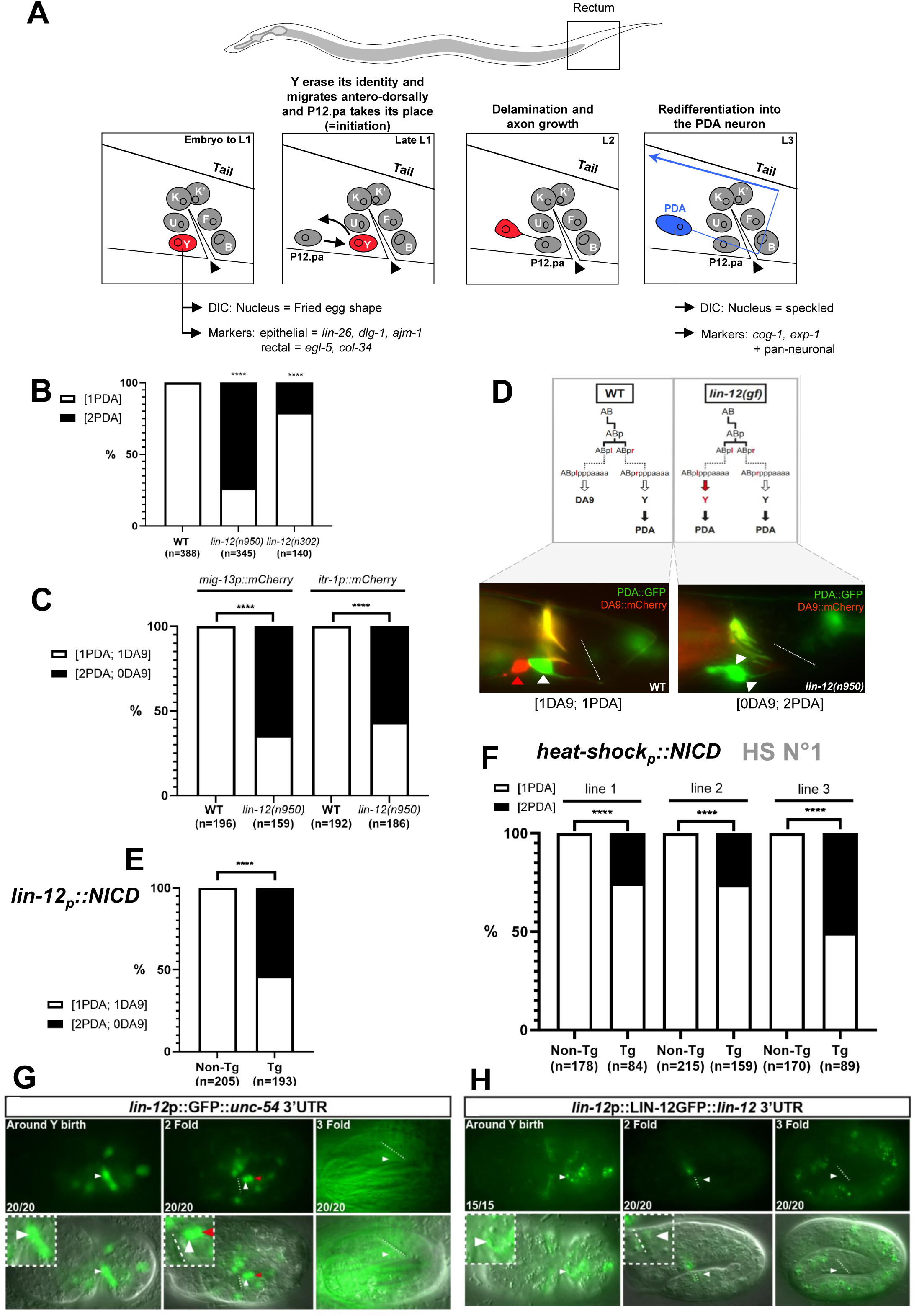
Ectopic Notch activity during early embryogenesis results in an extra transdifferentiation event. **A)** Timeline of the Y-to-PDA Td. In the L1 stage, the rectum of the worm is composed of 6 specialised epithelial cells (named Y, B, U, F, K and K’) arranged in three rings. These cells, as well as DA9^prog^, are born at the same time in the embryo around 290 min after the first cleavage^67^and are fully differentiated when the worm hatches. The Y cell is recognized by its position and fried-egg shaped nucleus using DIC, or by using epithelial or rectal markers. The Y cell keeps its epithelial identity until the end of the first larval stage, when Td is initiated. Y retracts from the rectum and migrates antero-dorsally while the P12.pa cell takes its place in the rectum to maintain the organ integrity. Concomitantly, the Y identity is erased in a dedifferentiation step *sensu strictu*, delaminates from the rectum. Later in L2 redifferentiation into a motoneuron begins, to adopt the “PDA” final identity by the L3 stage. The PDA neuron can be recognized by its position and its speckled nucleus using DIC or by using neuronal markers. This stereotyped event is identifiable and predictable in all WT animals. The rectal slit is indicating by the black arrowheads. **B)** Quantification (in %) of the 2 PDA phenotype in WT, strong *lin-12^Notch^* (*n950*) and mild *lin-12^Notch^*(*n302*) gain-of-function alleles with the *cog-1::gfp* PDA marker. **C)** Quantification (in %) of the “2 PDA and absence of DA9” phenotype in *lin-12^Notch^*(*n950*). The PDA and DA9 cells are visualized with *cog-1::gfp,* and *mig-13p::mCherry* or *itr-1p::mCherry* reporters respectively. **D)** DA9 and PDA in *lin-12^Notch^(gf)*. **Top panel**: Lineage tree and fate of ABplpppaaaa (DA9^prog^) and ABprpppaaaa (Y^prog^) cells in WT, and *lin-12^Notch^*(gf) animals. Dotted line represents cell division not displayed in the scheme (after Sulston et al. 1983). In WT worms, the ABprpppaaaa cell gives rise to the Y cell, which transdifferentiates to become a PDA neuron, while its contralateral ABplpppaaaa cell gives rise to the DA9 neuron. In *lin-12^Notch^(gf)*, both ABp/lpppaaaa cells give rise to Y cells, which are both competent to transdifferentiate, resulting in a “0 DA9; 2 PDA” phenotype in L4. **Bottom panel:** Pictures of the WT (left) and “2 PDA; 0 DA9” phenotype of *lin-12^Notch^(n950)* worms (right). The PDA and DA9 cells are visualized with *cog-1::gfp* and *itr-1p::mCherry* reporters respectively. White arrowheads: PDA neuron. Red arrowheads: DA9 neuron. **E)** Quantification (in %) of the number of PDA and DA9 neurons found in a *lin-12^Notch^p::*NICD*:: lin-12^Notch^3’UTR* transgenic line. Their morphology, position and the *cog-1::gfp* and *itr-1p::mCherry* molecular markers were used to identify the PDA and DA9 cells respectively. **F)** Quantification (in %) of worms exhibiting the “2 PDA” phenotype in three independent transgenic lines expressing the *hsp*::NICD construct, assessed with the *cog-1::gfp* reporter after a heat-shock during early-embryogenesis. **E-F)** n, total number of animals scored. Tg, transgenic animals. Non-Tg, non-transgenic (control) siblings. Data represent the mean of at least three biological replicates. Two-tailed P value is calculated using a Chi^2^ test. ****P < 0.0001, ***P < 0.001, ***P < 0.01, *P<0.05 **G)** Expression pattern and dynamics of *lin-12^Notch^p::gfp* (*rtEx727*). *lin-12^Notch^* is expressed in Y (arrowhead) during embryogenesis from its birth to the 2-fold stage. In the 3-fold stage, the diffuse GFP signal is due to expression of *myo-2p*::*gfp*, co-injection marker used to generate this transgenic line; no *lin-12^Notch^p::gfp* signal is detected in Y nor DA9 at this stage. **H)** Expression pattern and dynamics of the LIN-12^Notch^ protein (*arIs41*). LIN-12^Notch^ is present in Y during embryogenesis from Y birth to the 2-fold stage (where a faint signal can still be detected). **G-H)** Dotted line, rectal slit. Anterior is to the left and ventral to the bottom. Inserts represent blown-up rectal areas. White arrowheads: Y cell. Red arrowheads: DA9 neuron. Numbers represent the fraction of worms displaying this representative expression pattern over the total number of worms scored.

Here we show that Notch signalling not only specifies the Y cell as rectal but also endows this cell with the ability to change its identity. Strikingly, ectopic Notch signal is sufficient to create, *in vivo,* another cell with the potential to transdifferentiate. RNAi and expression analyses showed that the extra Td utilises the same molecular mechanisms as in the natural Y-to-PDA Td^31,32,34^ and highlighted a set of genes downstream of the Notch signal that are strictly necessary for Y cell reprogramming and not for Y rectal identity. By supplying prolonged or transient Notch signal to the Y cell at different time points, we demonstrated that seeing the Notch signal at the wrong time can block Y Td by re-enforcing its rectal identity. Thus, while a Notch signal is crucial to allow a cell to change its identity, it can exert a paradoxical action on the Y cell fate. Finally, the careful analysis of the characteristics of the supernumerary PDA neurons and the cells they originate from, showed that i) this ectopic Td requires a favourable cellular environment to occur; and ii) that Notch signal levels do not impact on when Y Td occurs. Together, our findings highlight that Notch signalling dynamics and tight regulation *in vivo* are essential to endow a unique cell with the capacity to transdifferentiate.

## Results

### Ectopic Notch signal is sufficient to induce an extra cell with the capacity to transdifferentiate

A second Y cell is found in the specific Notch gain-of-function (*gf*) allele, *lin-12^Notch^ (n137)*^17,28^. We had previously shown that this second Y cell transdifferentiates into a second PDA neuron later in development^28^ (Fig1A). To understand the role of Notch signalling in Y-to-PDA Td, we first investigated if the generation of more PDA neurons is observed in other *lin-12^Notch^(gf)* alleles. We found that all *lin-12^Notch^(gf)* alleles examined [i.e. *n950* or *n302*, strong and milder *lin-12^Notch^(gf)* alleles respectively] exhibited an extra PDA neuron (Fig1B). Concomitantly, a neuron named DA9 is also absent every time that a second PDA is found (Fig1C). In wild type (WT), after its final division, the ABplpppaaaa blastomere [hereafter termed DA9^prog^ to signify early undifferentiated DA9 cell] differentiates into the DA9 neuron. In *lin-12^Notch^(gf)* mutants, DA9^prog^ is mis-specified into a supernumerary Y cell^17,28^ that later transdifferentiates into a supernumerary PDA neuron^28^ (Fig1C-D). Together with the fact that extra Y cells made in other mutant backgrounds do not have the capacity to transdifferentiate into a PDA neuron^28^, our results suggest that Notch signalling not only promotes a rectal Y fate, but also confers the competence to transdifferentiate. By contrast, *lf* or *gf* alleles of *glp-1*, the second *C. elegans* Notch receptor gene, had no impact on the number of PDA neuron, indicating that GLP-1^Notch^ is likely not involved in Y Td (FigS1A).

We next examined if Notch activity is sufficient to obtain supernumerary PDA neurons. To this aim, we over-expressed an exogenous Notch IntraCellular Domain (NICD) protein, a constitutively active form of the LIN-12^Notch^ receptor, *in vivo*. First, NICD was expressed under the *lin-12^Notch^* promoter, provoking the activation of the Notch pathway only in the cells that normally express *lin-12^Notch^*. Strikingly, worms carrying the *lin-12^Notch^p::NICD* transgene had 2 PDA neurons, instead of 1 in WT, and an associated absence of the DA9 neuron, a phenotype similar to *lin-12^Notch^(gf)* alleles (Fig1E). Next, we investigated whether supernumerary Tds (i.e., more than one PDA) could also be obtained when we exogenously activated Notch ubiquitously. To this aim, a heat-shock (hs) promoter was used to drive ubiquitous expression of NICD in the worm. Remarkably, we found that a ubiquitous pulse of an exogenous Notch signal at the time of Y birth also results in the formation of an extra PDA, indicative of an extra Td (Fig1F). Therefore, an embryonic pulse of Notch signal is sufficient to later produce a supernumerary Td in live animals. Altogether, these results indicate that the Notch signal is sufficient to create cells competent to transdifferentiate, *in vivo*.

To further investigate when and where Notch signalling acts to promote a supernumerary Td, we determined the expression of *lin-12^Notch^* in the rectal area: a *lin-12^Notch^* transcriptional reporter^35^ is expressed in the ABprpppaaaa cell [hereafter called Y^prog^ before it has differentiated into the rectal Y cell], and in the DA9^prog^ cell, from their birth to approximately the 2-fold stage in WT embryos (Fig1G, FigS1B). After this point, no residual signal was detected in Y^prog^ or DA9^prog^, showing that *lin-12^Notch^*transcription takes place only during a restricted time window. Analysis of a *lin-12^Notch^* translational reporter, known to rescue *lin-12^Notch^ lf* mutants^36^, showed that the presence of the LIN-12^Notch^ receptor follows the exact same dynamic pattern as the *lin-12^Notch^* transcriptional reporter in the Y^prog^ cell (Fig1H, FigS1B), in accordance with previous reports that the LIN-12^Notch^ protein has a very short half-life^37^. Together with the phenotype of the *lin-12^Notch^(gf)* mutants, these results suggest that the LIN-12^Notch^ signal could act within the Y^prog^ or DA9^prog^ cells to endow them with the competence to transdifferentiate.

### The supernumerary Y-to-PDA Td in *lin-12^Notch^(gf)* requires the same plasticity factors as the endogenous Td

We have previously identified 5 nuclear factors as being necessary for the initiation of Y Td: the members of the NODE complex (*ceh-6/Oct*, *sem-4*/*Sall*, *egl-27*/*Mta1*), *sox-2/Sox-2,* and the downstream factor *egl-5/HOX*^28,32^. Does the Td of the supernumerary Y cell in *lin-12^Notch^(gf)* require these same plasticity factors as the endogenous Y-to-PDA Td? We thus examined if RNAi inactivation of these factors in a *lin-12^Notch^(gf)* genetic background led to suppression of the “2 PDA” phenotype. Our results showed that these factors are also necessary for the Td of the supernumerary Y cell into a PDA neuron, as RNAi against these genes led to a significant reduction of the “2 PDA” phenotype and the appearance of a “0 PDA” phenotype (Fig2A, FigS1C-D). In addition, an assorted “0 DA9” phenotype is observed: DA9^prog^ is mis-specified into a Y cell in *lin-12^Notch^(gf)* mutant (“0 DA9”), that cannot transdifferentiate in absence of these factors (“0 PDA”). Using the same assay, we also found that *unc-3*, a factor we have shown to mediate the subsequent redifferentiation step^31^ (Fig1A), appears to be necessary for the formation of the supernumerary PDA, as expected if the same mechanisms are at play (Fig2B, FigS1E).

**Figure 2:**
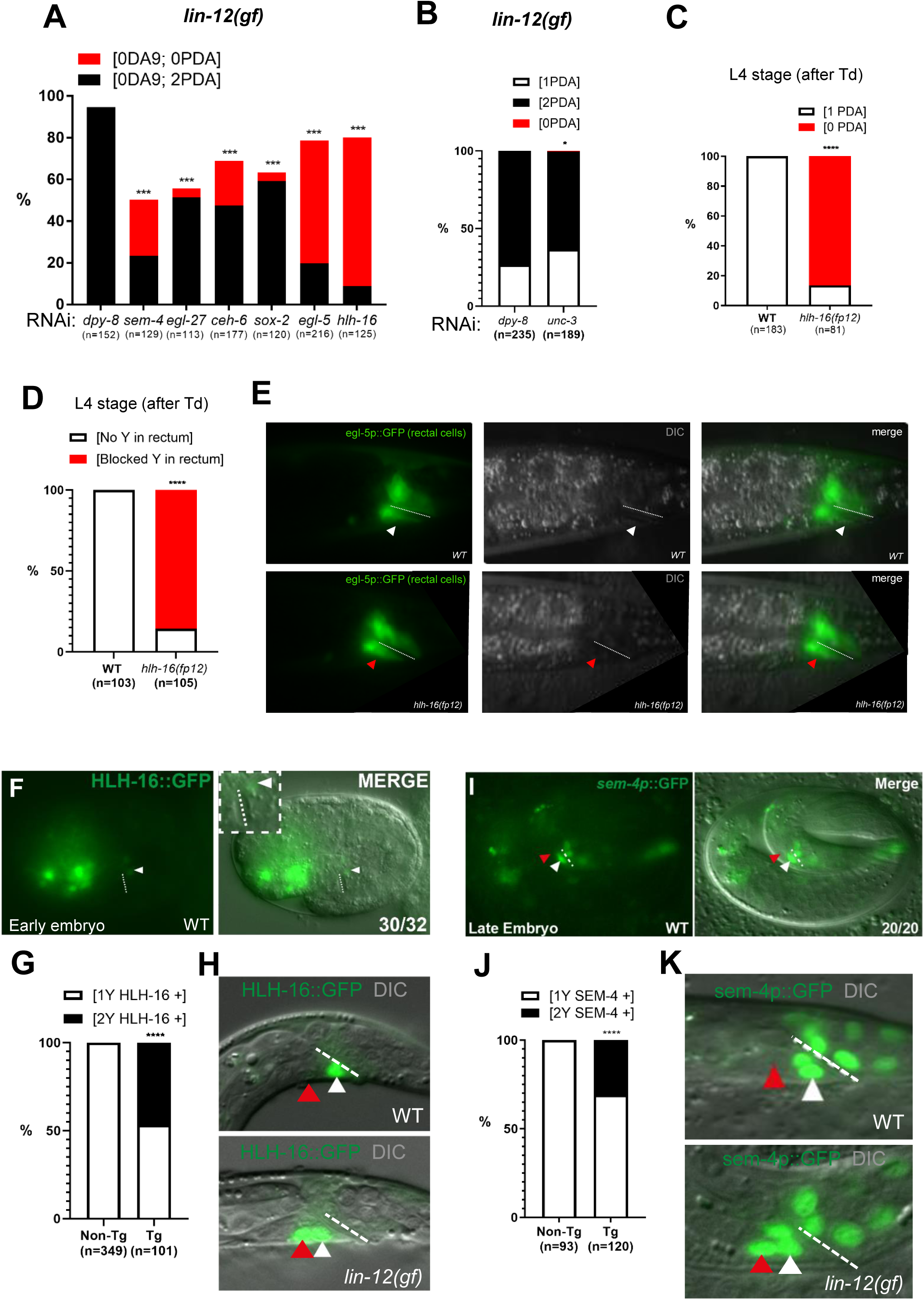
Supernumerary Td requires endogenous Td factors, unveiling *hlh-16* and *sem-4* as downstream of Notch. **A)** The plasticity factors necessary for the WT Y-to-PDA Td are also requires for the extra Td event (“2 PDA” phenotype) of *lin-12^Notch^(gf)*. The “2 PDA” phenotype (in %) is significantly suppressed when the plasticity factors (*sem-4, egl-27, ceh-6*, *sox-2, egl-5* and *hlh-16)* are knocked-down by RNAi in *lin-12^Notch^(n950)* (*dpy-8,* negative control). *cog-1::gfp* and *itr-1p::mCherry* were used as PDA and DA9 markers respectively. See also SIFig1B-C. **B)** Significant suppression of the *lin-12^Notch^(gf)* “2 PDA” phenotype (in %) when *unc-3* is knocked-down by RNAi in *lin-12^Notch^(n950). dpy-8,* negative control, *cog-1::gfp* as PDA marker. Of note, *unc-3*(RNAi) is poorly efficient and leads to a very low penetrance defect. **C)** Quantification (in %) of PDA presence at the L4 stage (after Td) in *hlh-16(fp12)* in comparison to WT as assessed with *cog-1* reporter. **D)** Quantification (in %) of a persistent Y cell in the rectum at the L4 stage (after Td) in *hlh-16(fp12)* in comparison to WT (no Y present anymore) as assessed with *egl-5* reporter. **E)** Picture of the phenotypes represented in (D). Top: WT L4 animals; bottom: *hlh-16(fp12)* L4 mutant. White and red arrowheads show the absence of Y in WT and a persistent Y cell in the rectum, respectively. Dotted line: rectal slit. Left panels, fluorescence microscopy; middle panels, DIC microscopy; right panels, merged. **F)** HLH-16::GFP is expressed in Y^prog^ but not in DA9^prog^ in the early WT embryo (1.5-fold). Insert represents blown-up rectal area. **G)** Quantification of the number of HLH-16+ Y cells in the rectum (in %) in WT and in *lin-12^Notch^p::NICD* L1 animals. **H)** HLH-16::GFP is expressed in Y WT L1 larva (white arrowhead). 2 Y cells expressing HLH-16::GFP (red and white arrowheads) are found in *lin-12^Notch^ p::NICD*. **I)** *sem-4p::gfp* is expressed in Y^prog^ but not in DA9^prog^ in the late WT embryo (3-fold). **F, I)** Numbers represent the fraction of worms displaying this representative expression pattern over the total number of worms scored in bottom right corner. **J)** Quantification of the number of SEM-4+ Y cells in the rectum (in %) in WT and in *lin-12^Notch^p::NICD* L1 animals. **K)** *sem-4p::GFP* is expressed in Y WT L1 larva (white arrowhead). 2 Y cells expressing *sem-4p::GFP* (red and white arrowheads) are found in *lin-12^Notch^p::NICD*. **A-D, G, J)** n, total number of animals scored. The % of each phenotype observed, indicated under brackets above, is represented. Data represent the mean of at least three biological replicates. Two-tailed P value is calculated using a Chi^2^ test. Tg: transgenic; non-Tg: non transgenic (control) siblings. ****P < 0.0001, ***P < 0.001, ***P < 0.01, *P<0.05; **E-F, H-I, K:** anterior to the left, and ventral at bottom. The dotted line represents the rectal slit.

During our initial genetic screen to identify the molecular players involved in Y-to-PDA Td, we retrieved the *fp12* allele, which potentially affects another player^30^. *fp12* had been mapped using EMS-density mapping^30^, which we confirmed by SNP mapping and found to affect the *hlh-16/Olig* gene (FigS2A), a nuclear bHLH protein^38^. *fp12* bears a point mutation (G →A) in the exon 2 splicing acceptor site, leading to a *lf* mutation. In *hlh-16(fp12)* mutant, a very high proportion of “0 PDA” is observed (Fig2C). Two experiments confirmed that *fp12* is a *hlh-16 lf* mutant: i) RNAi experiments against *hlh-16* phenocopied *hlh-16(fp12)* mutant (FigS1C) and ii) over-expression of WT *hlh-16* under a *hlh-16* promoter in the *hlh-16(fp12)* mutant background rescued the “0 PDA” phenotype (FigS2B). Further analyses revealed that, in *hlh-16(fp12)* mutant, the Y cell is born and differentiates into a rectal cell as in the WT (FigS2C). However, it then remains rectal at its original position, with its characteristic morphology and expressing the *egl-5* rectal marker. Td is never initiated in the L2 stage (Fig2D-E). These results suggest that *hlh-16/Olig* is implicated in Y Td initiation, rather than in Y cell fate determination.

To confirm that *hlh-16* is required for the Td initiation as opposed to the earlier Y differentiation into a rectal cell, we examined if late over-expression of *hlh-16* would rescue the Td defect seen in *hlh-16(fp12)* mutant. Late *hlh-16* over-expression, using the *col-34* promoter^32^, rescued the “0 PDA” mutant phenotype (FigS2B). Thus, *hlh-16* is necessary for Y cell Td at the time of its initiation (and not at its birth), in accordance with recent findings by Rashid et al^39^. In addition, combined with our findings that *hlh-16* is only expressed in the Y cell among the rectal cells (Fig2F-H), these results strongly suggest that *hlh-16* acts in a cell-autonomous manner.

Interestingly, *hlh-16* acts at the same step as the plasticity cassette factors necessary for the initiation of Y Td. We therefore examined if *hlh-16* was also necessary for the *lin-12^Notch^*-induced supernumerary Td. RNAi inactivation in *lin-12^Notch^(gf)* mutant showed that *hlh-16* is required for both the supernumerary and the endogenous Y-to-PDA Tds (Fig2A, FigS1C-D).

Altogether, our results suggest that the Notch-induced ectopic Td event requires the same molecular mechanisms as the endogenous Y-to-PDA Td.

### HLH-16 and SEM-4, two key factors for Td initiation, are downstream of Notch signal

We next asked if the Notch pathway conferred competence to transdifferentiate by promoting plasticity factors expression. To this aim, we compared expression of the plasticity factors (NODE complex (*ceh-6/Oct*, *sem-4*/*Sall*, *egl-27*/*Mta1*), *sox-2/Sox-2, egl-5/HOX* and *hlh-16/Olig*) in WT and worms carrying the *lin-12^Notch^p::NICD* transgene (active form of the *lin-12^Notch^*receptor).

We found two types of patterns: i) genes endogenously expressed in both the Y^prog^ and DA9^prog^ cells in WT animals since their birth. This is the case for the *egl-5*, *sox-2* and *ceh-6* genes (FigS2D-F). Since these genes are already expressed in the DA9^prog^ cell since its birth in absence of ectopic Notch, we concluded that these genes are not downstream of Notch. As *egl-5, sox-2* and *ceh-6* genes are necessary for Y Td, but not Y identity^28,34^, these genes could either contribute to a Notch-permissive context, or allow the Y cell Td independently of Notch. And, ii) genes that, besides being naturally expressed in the Y^prog^ cell, are only observed in the extra Y cell coming from the mis-specification of DA9^prog^ after receiving an exogenous *lin-12^Notch^* signal. This is the case for the *hlh-16* and *sem-4* genes (see the Y cell and a second cell persistently expressing *hlh-16* or *sem-4* in Fig2G-H and Fig2J-K). A second Y cell is observed in *lin-12^Notch^(gf)* mutant worms with the same penetrance as the “2 PDA” phenotype. Consistently, *hlh-16* or *sem-4* expression starts in the Y cell quite some time after its birth (Fig2F-I). Indeed, *sem-4* is observed in Y rather late (when the embryo is fully elongated, Fig2I), while *hlh-16* starts to be expressed in comma stage embryos (Fig2F), and their expression is maintained in the Y cell during the L1 stage (Fig2G-K). It thus appears that in endogenous Y^prog^ and in DA9^prog^ experiencing an ectopic Notch signal, expression of *hlh-16* and *sem-4* is triggered downstream of the Notch signal. Of note, expression of *hlh-16* and *sem-4* persists until Td initiation, while LIN-12^Notch^ expression ceases during embryogenesis (2-fold, Fig1G-H), suggesting that the Notch signal is necessary to promote their initial expression but that expression maintenance is most likely Notch-independent.

Therefore, the Notch pathway enables the Y cell competence to transdifferentiate by promoting plasticity factors expression such as *hlh-16* and *sem-4*.

### A permissive context for Notch signal is necessary to induce a cell with the capacity to transdifferentiate

Interestingly, when ubiquitously expressing NICD under the hs promoter in the whole animal, we found that these animals had two, and not many, PDA neurons (Fig1H). In particular, the other rectal cells, born at the same time as Y^prog^, do not appear to change their identity nor their typical fried-egg-shaped nuclear morphology. We next tested if using a promoter driving robust and maybe longer NICD expression specifically in these cells could allow their conversion into supernumerary PDA neurons. To this end, the constitutively active NICD was expressed under a rectal promoter, *egl-5p*^28^, which expression starts shortly after the rectal cells’ birth (FigS3A). The identity of the rectal cells was monitored at the L4 stage using both appearance under DIC microscopy and rectal markers (*lin-26*, *lin-48* and *mab-9,* Fig3A-C). We found that a continuous Notch signal in these cells did not result in a switch in their identity, and no extra PDA neurons were observed (Fig3D). These other rectal cells, therefore, represent a refractory context for the Notch-promoted Td, highlighting the importance of a particular cellular context for the efficiency of a given reprogramming cue. Thus, the Notch signal, which is sufficient to endow a cell with the competence to transdifferentiate, does so in a permissive context.

**Figure 3:**
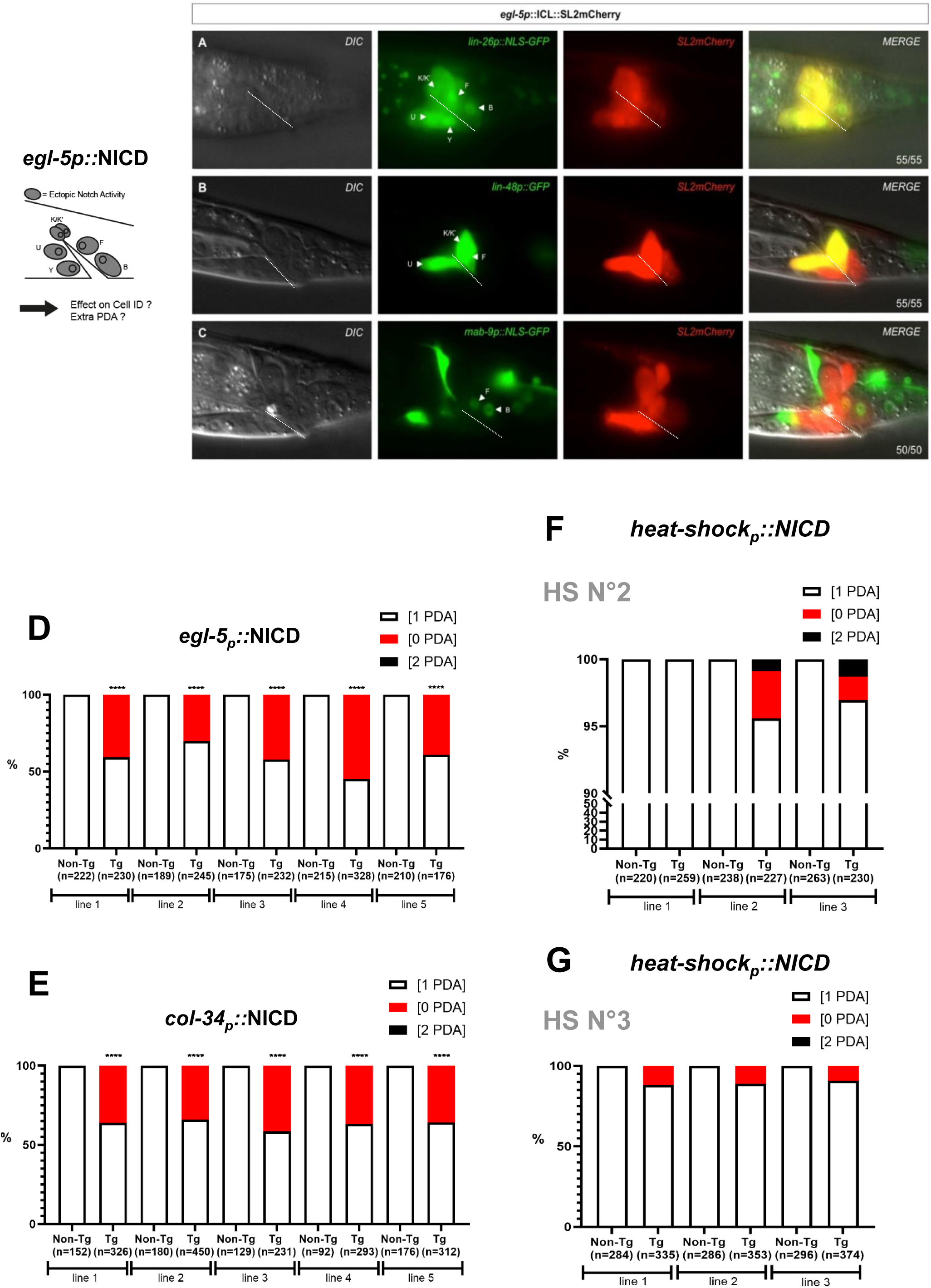
Exogenous Notch signal in other cells than DA9^prog^ is not sufficient to provoke a <u>change of identity</u>. Left, experimental approach: overexpression of the NICD construct in all the rectal cells using the *egl-5* promoter. **A-C)** Representative pictures of the specified marker expression in a L4 worm (A, C) or a L3 worm (B), showing that the rectal identity of the B, U, F, K and K’ rectal cells is not affected by NICD overexpression. SL2::mCherry shows NICD expression in the rectal cells. No constipation is observed, showing that these rectal cells are fully functional. Bottom right corner, number of animals showing the represented phenotype over the total number of animals scored. Anterior is to the left and ventral to the bottom. Arrowheads show the position of the designated cells. Dotted line, rectal slit. **D-E)** High penetrance of Td defect [0 PDA] (in %) is found in transgenic (“Tg”) worms expressing *egl-5p*::*NICD::SL2mCherry* (**D**) or *col-34p::NICDGFP* (**E**) in five independent transgenic lines. No extra Td [2 PDA] was observed. Non-transgenic (“Non-Tg”) control siblings are WT as observed with *cog-1::gfp* reporter. See SIFig3A for the timing of *egl-5* and *col-34* expression. **F-G)** Quantification (in %) of worms exhibiting an extra Td [2PDA] or no Td [0 PDA] phenotype after a heat-shock during (**F**) mid-embryogenesis (HS 2, SIFig3A) and (**G**) before Td initiation in late L1 stage (HS 3, SIFig3A), in three independent transgenic lines expressing the *hsp*::NICD construct, as assessed with *cog-1::gfp* reporter. **D-G)** n, total number of animals scored. Data represent the mean of at least three biological replicates. Two-tailed P value is calculated using a Chi^2^ test. ****P < 0.0001, ***P < 0.001, ***P < 0.01, *P<0.05

### Dichotomic action of Notch signalling on Y-to-PDA Td

A closer look at the worms carrying the transgene *egl-5p::NICD* showed to our surprise that a large number of them did not have a PDA neuron (“0 PDA”, Fig3D). In addition, the rectal Y cell was still present in the rectum at the L4 stage (Fig3A), suggesting that this treatment resulted in a block of the endogenous Td of the Y cell. We therefore analysed how an exogenous Notch signal can both promote formation of ectopic PDAs or lead to a Td blockage (“0 PDA” phenotype), hereafter called “dichotomic effect”. The *egl-5* promoter used to express NICD in the rectal cells is active from Y birth to the initiation of Td (FigS3A). However, in WT worms, Notch receptor expression stops after the 2-fold stage (Fig1G-H, FigS3A). This suggests that the “No Td” (“0 PDA”) phenotype obtained when expressing NICD under the *egl-5* promoter could be a consequence of a prolonged Notch signal in the Y cell. To determine if the timing of the exogenous Notch signal was crucial for this dichotomous effect on Y Td, we expressed NICD in the rectal cells later, starting from the late embryonic 3-fold stage, using the *col-34* promoter^32^ (FigS3A). This also resulted in a strongly penetrant “0 PDA” phenotype (Fig3E), suggesting that it is either a continuous or a late Notch signal that was detrimental to Y Td. We next tested the impact of the Notch signal at specific time points (FigS3A) by providing a pulse of NICD under the hs promoter: i) at the embryonic 2-fold stage, when endogenous *lin-12^Notch^*expression disappears (see HS 2, Fig3F), and ii) during the L1 larval stage, just before the initiation of the Td process in WT worms (see HS 3, Fig3G). While the expression of NICD at the 2-fold stage resulted in a few instances of the “0 PDA” phenotype (and also in few “2 PDA”, Fig3F), expression at the time of Td initiation resulted in a significant number of “0 PDA” instances, and an absence of the “2 PDA” phenotype (Fig3G). These results suggest that: i) the Notch pathway could not promote the conversion of DA9^prog^ into an extra competent Y cell anymore after the 2-fold stage (no “2 PDA” phenotype); ii) the closer the Y cell is to Td initiation when it encounters ectopic Notch pathway activation, the stronger the “0 PDA” phenotype is. Thus, the embryonic 2-fold stage represents a critical check point for the Notch signal to be interpreted as a pro-or anti-Td signal (mix of “0 PDA” and “2 PDA” phenotypes). In addition, while the different drivers used may lead to variable levels of the Notch signal, our data suggest that the critical parameter is encountering the Notch signal at the Td initiation time, independent of Notch levels. Altogether, these results suggest that the time window during which the Y cell receives a Notch signal is key for its capacity to transdifferentiate; seeing Notch signalling, even transiently, at the time of the Td initiation blocks the identity conversion.

We then examined if late, ectopic Notch activation blocked Y Td cell-autonomously, or if it was due to Notch activation in (an)other cell(s). With a particular focus on the rectal cells, we drove the expression of NICD using various promoters active in subsets of them and examined the presence of PDA. First, we used the *lin-48* promoter, which is only active in the F, U, K and K’ rectal cells^40,41^ (Fig4A). Such partial rectal expression does not result in a “0 PDA” phenotype (Fig4B), implying that an activated Notch signal in F, U, K and K’ is not responsible for the Td defect observed with the *egl-5p::NICD* transgene. This suggested that the Notch signal acts either in the Y or the B cell. To discriminate between the B and Y cells, we used the mosaic expression our *egl-5* transgene. We focused on the expression of the transgene in the B cell and found no correlation between NICD expression in the B cell and the appearance of a “0 PDA” phenotype (Fig4C). We also used the *egl-20p* promoter, previously described as being expressed in all rectal cells except Y^42^. We however observed, similarly to others^43^, that *egl-20* is in fact expressed in Y with a high degree of mosaicism (FigS3B). Nevertheless, most of the worms displaying a “0 PDA” phenotype did not express NICD in the B cell (Fig4D*-*FigS3C), further suggesting that it is Notch activation in the Y cell that blocks its Td.

**Figure 4:**
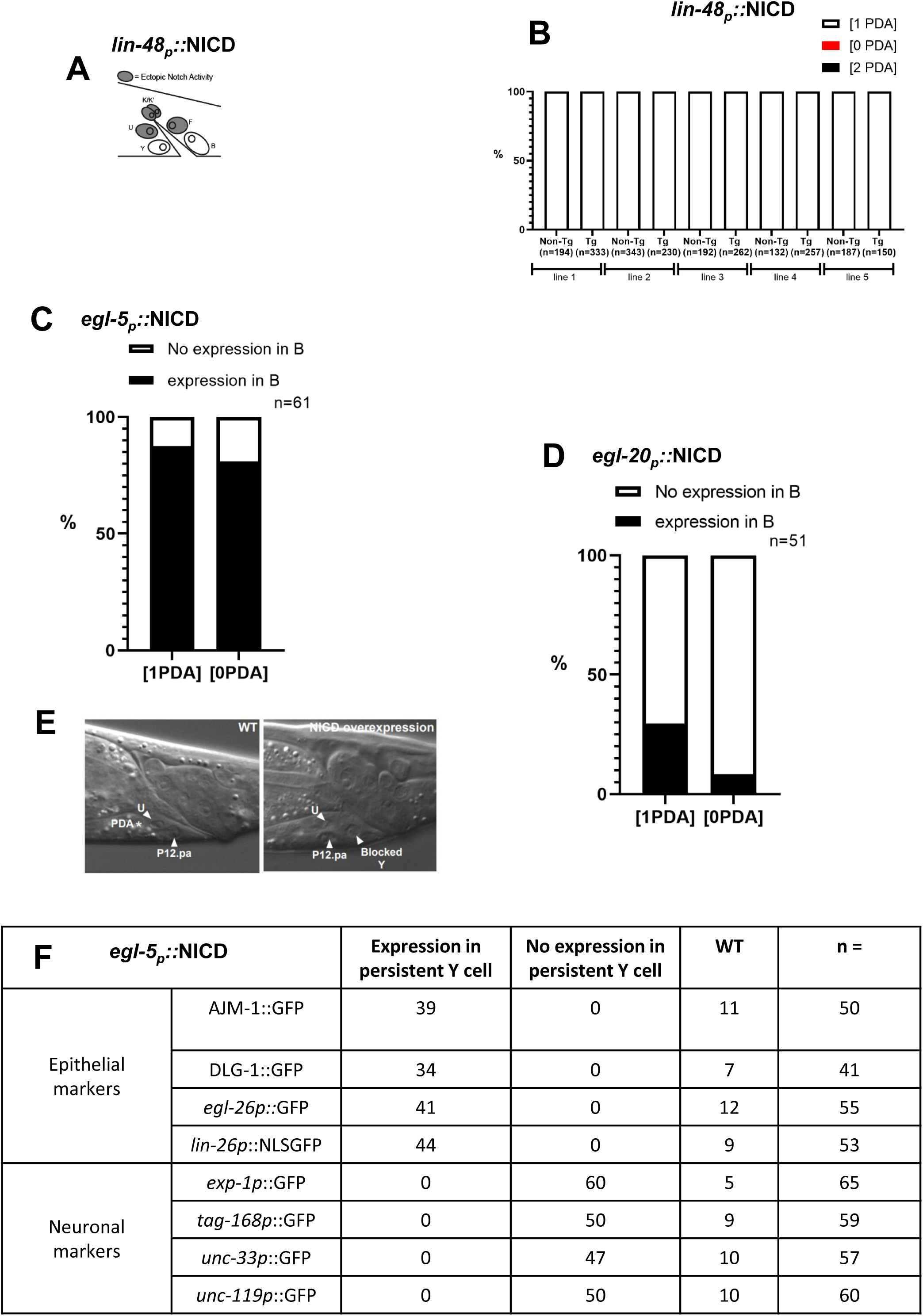
A late Notch signal cell-autonomously blocks initiation of Td. **A)** Experimental approach: overexpression of the NICD construct in a subset of the rectal cells using the *lin-48* promoter. **B)** No Td defects ([0 PDA], in %) is observed when NICD (*lin-48p*::*NICD::SL2mCherry)* is expressed in the F, U, K and K’ rectal cells in five independent transgenic lines as assessed using *cog-1::gfp* reporter. Tg, transgenic worms; Non-Tg, non-transgenic WT siblings; n, total number of worms scored. **C-D)** A similar proportion (**C**) or even a higher proportion (**D**) of animals (in %) exhibit a Td defect [0 PDA] when NICD is not expressed (white bar) in the B cell using a mosaic *egl-5* driver (**C**) or a mosaic *egl-20* driver (**D**), revealing no correlation between Td defect and expression in B; total number of animals scored = 61 (C) and 51 (D). **E)** Representative DIC picture of the persistent Y cell in L4 transgenic animals expressing ectopic activated Notch in the rectal cells (*egl-5p::NICD*). Left, WT worm where Y has turned into a PDA (star). Right, L4 worm with ectopic Notch activation in Y. The Y cell (arrowhead) is blocked before Td initiation and found at its original position in the rectum; the rectal U and P12.pa cells (arrowheads) are also visible. Anterior is right and ventral is to the bottom. **F)** The persistent Y cell remains rectal. Epithelial and neuronal markers in the persistent Y cell (expressed in number of animals) in worms expressing an integrated NICD construct under the *egl-5* promoter (*egl-5p::NICD*). When Y Td is blocked, a persistent Y cell is found at its original position. This Y cell remains rectal. See also FigS4. n, total number of animals scored.

Altogether, these data suggest that Y Td is blocked by Notch over-activation in a cell-autonomous manner.

### Late Notch at the time of Td initiation blocks it by over-imposing a rectal identity

Why does a late Notch signal lead to an impairment of Y Td? To address this question, we examined the specific step at which Y Td is blocked, by analysing the morphology and position of the Y cell, plus the markers it expresses when it receives a late Notch signal. In WT worms, after Td has taken place, only two hypodermal cells are present in the anterior part of the rectum (P12.pa and U, Fig 4E). Interestingly, in worms overexpressing the NICD transgene, three hypodermal-like cells are present in the anterior part of the rectum, including the Y cell (Fig 4E). This mimics the phenotype of mutants where Td is blocked before initiation and the Y cell remains rectal at its original position^28,30–33^. In such mutants, the Y cell keeps its epithelial identity and does not express any neuronal markers. To investigate whether Y remains epithelial when exposed to a Notch signal late, we examined epithelial or rectal specific markers in worms expressing *egl-5p::NICD*. We used two epithelial junction markers, *AJM-1::GFP*, *DLG-1::GFP,* a rectal marker, *egl-26p::GFP*, and an epithelial differentiation marker, *lin-26p*::*NLS-GFP*^28^ (Fig4F-FigS4A-D). By contrast to the WT pattern, each of these markers remained expressed in the Y cell past the L2 stage, indicating that late Notch activity in Y leads to an early block of the Td process, with no retraction of Y from the rectum (Fig1A) and maintenance of its rectal and epithelial identity. To rule out the possibility that Y is blocked in a mixed identity state in which both epithelial and neuronal markers are expressed - something that never happens during the wild-type process^31^ - we looked for the potential expression of a second PDA marker, *exp-1p::GFP*, and three pan-neuronal markers, *tag-168p::GFP*, *unc-33p::GFP* and *unc-119p::GFP*, in the persistent Y. None of these were observed in the blocked Y (Fig4F, FigS4E-H). Collectively, our data show that late over-activation of the Notch pathway in the Y cell immediately prior to Td blocks its initiation by re-enforcing a rectal identity.

### Regulation of Notch transcription ensures that no endogenous Notch signal exists in the Y cell by the end of embryogenesis

Our discovery, that the time window during which the Y cell receives a Notch signal is crucial for proper Td, led us to examine how endogenous Notch activation is regulated. We first sought to identify the ligand(s) that activate it. Functional (3) or predicted (12) ligands and co-ligands exist in *C. elegans*^21,37,44^ and were tested for a role in Y-to-PDA using available mutants and RNAi in sensitised backgrounds (FigS5A-B). We found that the loss of *apx-1* or *lag-2* led to a Y-to-PDA defective Td (“0 PDA”), consistent with their role as ligands (Fig5A-B), while loss of any of the other potential ligands and co-ligands did not impair Td (FigS5B). Among the various strategies to enable fine control of Notch signal, ligand unavailability^45^ could result in switching off *lin-12^Notch^* receptor activity. Examination of the *lag-2* expression pattern showed that it is expressed from Y birth until the beginning of Td in the rectal B cell, which directly contacts and forms adherens junctions with Y^28^ (Fig5C). *apx-1* is also expressed in four cells close to the rectum at Y birth. After the embryonic 2-fold stage, two of these cells maintain this expression and remain in close contact with the rectum until at least beginning of Td (Fig5D). Thus, these two ligands are expressed around the Y cell, and their expression persists from its birth until at least the initiation of Td, a profile suggesting that the disappearance of the Notch signal in Y after the embryonic 2-fold stage is not due to the unavailability of the ligands. We also note that absence of ligand expression in the DA9^prog^ and Y^prog^ themselves, together with previous laser ablations^29^, indicate that a Notch inductive signal is involved in potentiating Y with the competence to Td.

**Figure 5:**
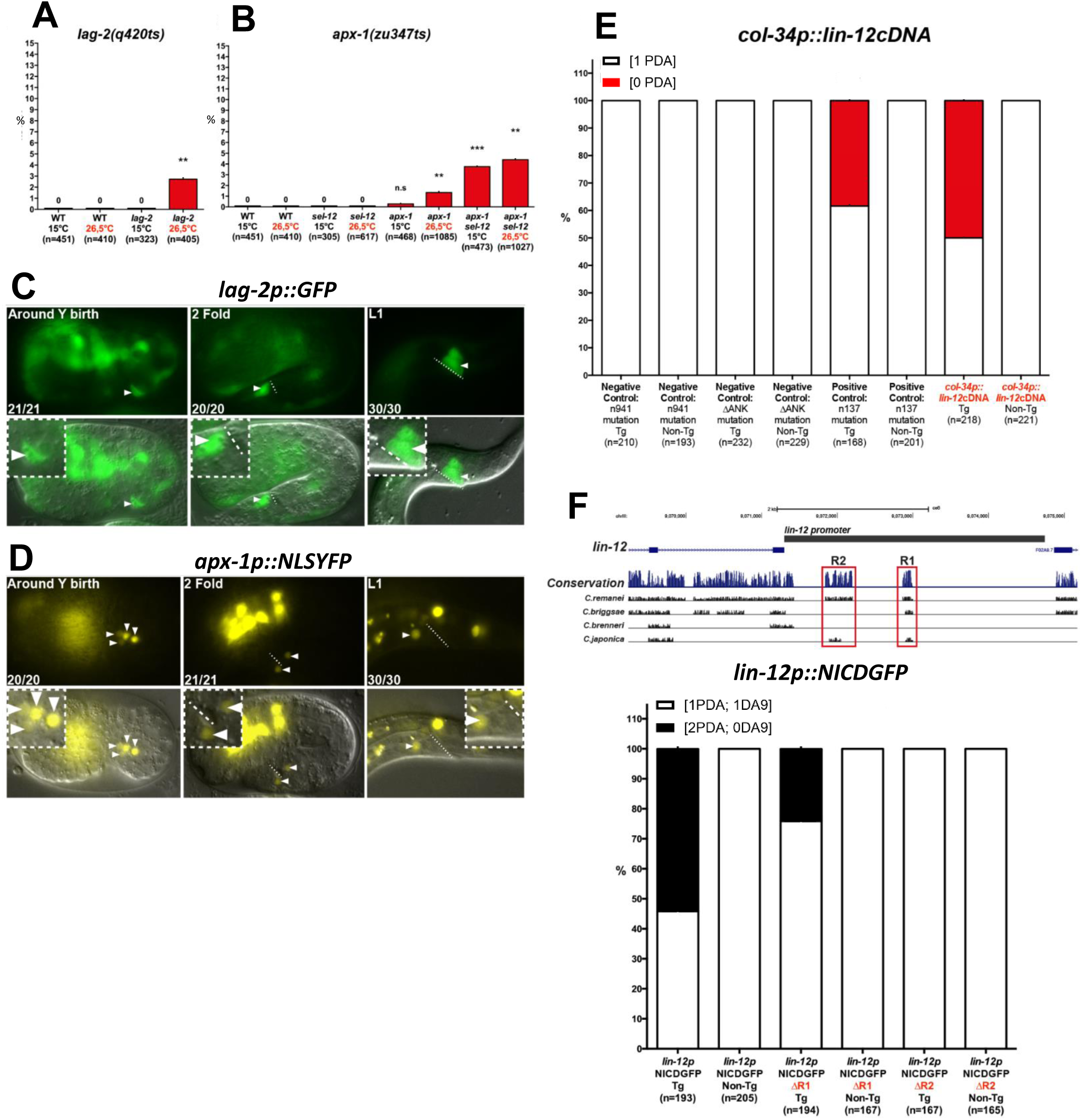
Regulation of Notch signal duration is achieved through transcriptional regulation A-D) *lag-2* and *apx-1* both act as ligands for activation of Notch in the Y cell. Permissive temperature, 15°C; restrictive temperature, 26,5°C. **A)** Quantification (in %) of a No Td [0 PDA] phenotype in *lag-2(q420ts)* mutant as assessed using the *cog-1::gfp* marker. **B)** Quantification (in %) of a No Td [0 PDA] phenotype in *apx-1(zu347ts)* mutant, alone or associated with *sel-12(ar171)* mutation. *sel-12* is one of the 2 presenilins in the worm and is an important component of Notch signal transduction. This allele does not exhibit any Y-to-PDA defect and is thus used as a sensitised background. **C-D)** Expression pattern and dynamics of *lag-2 and apx-1*. (**C**) *lag-2* is expressed in B (arrowheads) from Y birth until initiation of Td, after which it disappears from the rectal area. (**D**) Expression pattern and dynamics of *apx-1*. *apx-1* is expressed in four cells (arrowheads) close to the rectum during embryogenesis and at least 2 cells maintain expression throughout larval stages. Dotted line, rectal slit. Inserts represent blown-up rectal areas. Numbers represent the fraction of worms displaying this representative expression pattern over the total number of worms scored. **E)** Ligands are still available and functional when *lin-12^Notch^* is down-regulated. Last 2 bars, proportion of phenotypically WT or [0 PDA] worms carrying a *col-34p*:: *lin-12^Notch^(WT)cDNA* transgene, showing that functional ligands are available and activate a Notch receptor ectopically expressed after the embryonic 2-fold stage in the Y cell. Negative control transgenic lines: different inactive LIN-12^Notch^ receptors driven by *col-34p* [*col-34p::lin-12^Notch^cDNA(n941)* and *col-34p*:: *lin-12^Notch^cDNA(ΔANK)*]. Positive control: constitutively active full-length receptor driven by *col-34p* [*col-34p::lin-12^Notch^cDNA(n137)*]. As predicted, worms expressing inactive LIN-12^Notch^ receptors exhibit a WT Td, while a No Td [0 PDA] phenotype is found in transgenic (Tg) positive control worms, compared to non-transgenic (non-Tg) siblings. Only one transgenic line is depicted here; additional lines are depicted in FigS6A-D. **F)** Upper panel: the *lin-12^Notch^* promoter region (horizontal black rectangle) used to drive NICD expression. The blue track represents the overall conservation of the DNA sequence between different *Caenorhabditis* species. The black tracks represent the conservation of the DNA sequence of interest between one particular species and *C. elegans* (UCSC genome browser). The red boxes identify two conserved regions (R1, R2) of the *lin-12^Notch^* promoter further tested. Bottom panel: Quantification (in %) of the number of PDA and DA9 neurons found in transgenic lines expressing NICD under different *lin-12^Notch^* promoter variants. First 2 bars: *lin-12^Notch^p*:*:NICD* transgenic line. A [2 PDA] (and a concomitant [0 DA9]) phenotype appears in transgenic animals, while all the non-transgenics sibling are WT (same data as Fig1E). Middle 2 bars: *lin-12^Notch^p*(ΔR1)*::NICD* line. Deletion of the R1 region does not abolish transgene activity suggesting a minor role in the control of *lin-12^Notch^* timely expression. Left-most 2 bars: *lin-12^Notch^p*(ΔR2)*::NICDGFP* line. Deletion of the R2 region leads to absence of the [2 PDA] phenotype and all transgenic worms are WT, indicating that transgene expression has been affected. Additional transgenic lines related to this figure are depicted in SI6E-G. Tg, transgenic worms; non-Tg, non-transgenic siblings. **A-B, E-F)** n, total number of animals scored. Data represent the mean of at least three biological replicates. Two-tailed P value is calculated using a Chi^2^ test. ****P < 0.0001, ***P < 0.001, ***P < 0.01, *P<0.05

Are the ligands still functional after LIN-12^Notch^ has disappeared from Y? If yes, they should be able to activate an ectopic full-length LIN-12^Notch^ receptor re-expressed in Y late, after the endogenous one had disappeared (using a *col-34* promoter), and thus block Td. We indeed observed a No Td [0 PDA] defect, while non-transgenic controls were WT (Fig5E, FigS6A). These results show that the ectopic LIN-12^Notch^ receptor has been activated by the surrounding ligands, confirming that its ligands are indeed not only available but also functional until initiation of Td. As negative controls, we used forms of the LIN-12^Notch^ receptor that cannot be activated: a receptor-dead form [*lin-12^Notch^*cDNA bearing the premature stop codon in the extracellular EGF repeats as found in the null allele *lin-12^Notch^(n914)*^47^] and a LIN-12^Notch^ receptor unable to signal [*lin-12^Notch^*cDNA bearing a deletion of the seven ankyrin repeats of the intracellular part of LIN-12^Notch^ receptor^46^]. No inhibition of Y-to-PDA Td were observed in transgenic animals expressing these two constructs in Y late, as expected (Fig5F, FigS6B-C), suggesting no ectopic activation of the Notch pathway. As a positive control, a full length mutant receptor that is constitutively active was expressed [*lin-12^Notch^* cDNA containing the mutation found in the *gf lin-12^Notch^(n137)* allele, which has been shown to induce ligand-independent receptor activation^44,47,48^]. Transgenic animals expressing this construct exhibited a penetrant “0 PDA” phenotype (Fig5E, FigS6D), validating our conclusions.

Several lines of evidence point to a regulation of *lin-12^Notch^* expression over time in the Y cell: i) the LIN-12^Notch^ receptor disappears from the Y cell after the 2-fold stage (Fig1F); ii) the transcriptional *lin-12^Notch^* reporter showed the same expression dynamics (FigIE); since the transcriptional reporter lacks the *lin-12^Notch^* 3’UTR, these data suggested a transcriptional - rather than post-transcriptional - regulation of *lin-12^Notch^* ^35,36^(FigS4C). To test if the promoter of *lin-12^Notch^* is the main effector of LIN-12^Notch^ activity regulation, we re-examined our results expressing the NICD fragment under control of the *lin-12^Notch^* promoter (Fig1E). If *lin-12^Notch^* promoter is sufficient to recapitulate the brief WT pulse of Notch expression in the embryo, NICD expression under *lin-12^Notch^*promoter should lead to activation of the Notch pathway in Y^prog^ and DA9^prog^, and induce the appearance of a supernumerary Td [2 PDA]. If not, NICD would be expressed after the embryonic 2-fold stage in Y, leading to a block of Td [0 PDA]. Worms carrying this transgene displayed a “2 PDA” phenotype while the controls were all WT (“1 PDA”) (Fig1E,5F, FigS6E). These results show that down-regulation of the Notch signal in the Y cell is achieved through transcriptional modulation during embryogenesis, a crucial checkpoint to allow the subsequent initiation of Y Td.

We further identified two highly conserved regions (R1 and R2) between different *Caenorhabditis* species in the *lin-12^Notch^*promoter (Fig5F), with deletion of the R2 region completely abolishing the activity of the transgene (Fig5F, FigS6F-G). In conclusion, we have found that endogenous Notch expression is tightly regulated at the transcriptional level during embryogenesis, and a conserved region in its promoter is necessary for its transient expression during Y Td.

### Notch levels do not regulate the timing of Y-to-PDA Td

The dichotomic effect of Notch on the Y-to-PDA Td could suggest an involvement of the Notch pathway in regulating the timing of this reprogramming event. We were particularly intrigued by the suggestion that Notch levels modify the temporality of Y-to-PDA Td, as recently proposed in Rashid et al^39^. They reported premature occurrences of Y-to-PDA Td at the onset of the L1 stage instead of the beginning of the L2 stage in WT, in a specific *lf* allele of *lin-12^Notch^, n676n930,* that has been described as a weak *gf* allele in some particular contexts at 15°C^49^. Indeed, using 15°C as a putative *gf* temperature, they reported that the Y cell already exhibited an axon at the beginning of the L1 stage, while still expressing genes observed in the rectal Y cell (*ngn-1* and *hlh-16*), leading to the designation of this cell as a “precocious PDA”. However, this putative “precocious PDA” then failed to develop into a complete PDA, as suggested by the absence of PDA markers expression in older L4 animals^39^. Given their conclusion that a weak *lin-12^Notch^ gf* activity appeared to provoke an earlier Y cell reprogramming, they posited that the level of Notch signal regulates the timing of the Y-to-PDA Td.

However, we had previously reported that this specific *lin-12^Notch^*(*n676n930*) allele behaves as a *lf* at 15°C in the Y cell, resulting in no Y cell, hence 0 PDA, made^28^. Indeed, besides its role in the Y cell’s competence to transdifferentiate^28^, the Notch pathway is implicated in cell fate decision between a DA9 neuron and a Y rectal cell fate^17^. Specifically, Notch signalling has been shown to be necessary and sufficient for the Y cell fate^17^. In presence of high Notch levels Y^prog^ becomes rectal while no or low Notch levels lead to mis-specification as a DA9 neuron; both fates appear independent of each other, as shown by ablation experiments^50^. Consequently, in Notch *lf* alleles, such as *lin-12^Notch^(n676n930)* at 25°C and at 15°C, Y^prog^ is mis-specified as an extra DA9 neuron instead of a Y cell^28^. We thus re-examined this issue. Consistent with our previous report, we found that growing *lin-12^Notch^(n676n930)* at 15°C results in an absence of PDA, as would be expected for *lf* activity (Fig6A). Additionally, we never observed the distinct phenotype of *lin-12^Notch^(gf)* alleles (“2 PDA” and an absence of DA9, Fig1C-E) in *lin-12^Notch^(n676n930)* raised at 15°C. We confirmed by sequencing the presence of the *lin-12^Notch^ (n676n930)* mutations in our strain, excluding a *lin-12^Notch^* genetic issue as the basis for our diverging results from Rashid et al^39^.

**Figure 6:**
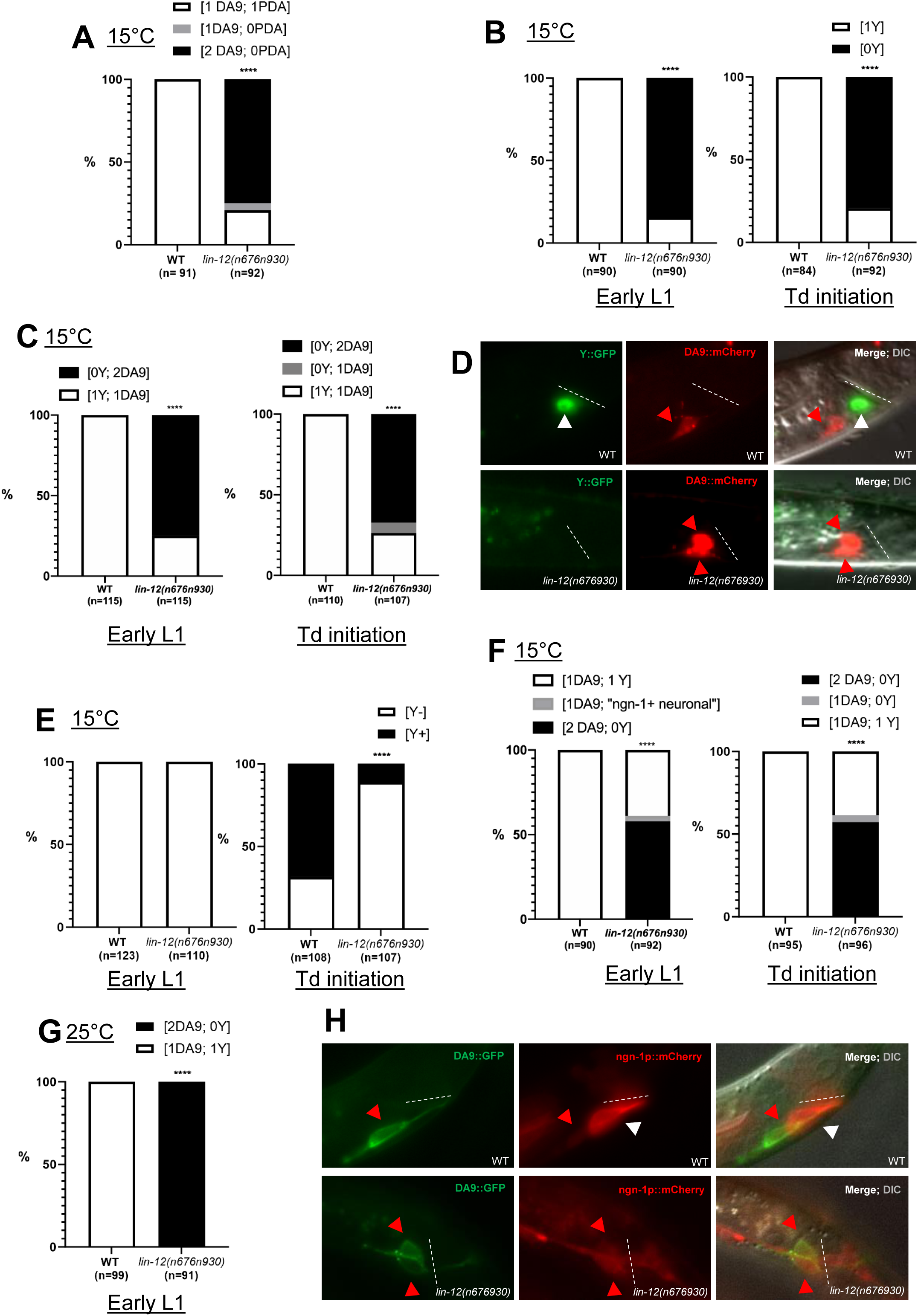
Notch level regulates the cell fate decision between DA9 and Y/PDA fates. **A)** Quantification (in %) of PDA and DA9 assessed with respectively *cog-1* and *itr-1* reporters in *lin-12^Notch^(n676n930)* at 15°C. **B)** Quantification (in %) of Y cell in *lin-12^Notch^(n676n930)* at 15°C assessed with *egl-5* reporter at early L1 stage (left) and early L2 stage (right). [1Y], WT Y cell (without any axon). **C)** Quantification (in %) of Y and DA9 cell assessed with respectively *hlh-16* and *mig-13* reporters in *lin-12^Notch^(n676n930)* at 15°C at early L1 stage (left) and early L2 stage (right). **D)** Representative picture of [1Y; 1DA9] WT phenotype and [0Y; 2DA9] observed in *lin-12^Notch^(n676n930)* L1 animals at 15°C, assessed with respectively *hlh-16* (Y::GFP) and *mig-13* (DA9::mCherry) reporters. **E)** Quantification (in %) of Y cell expressing *ngn-1(dev137)* endogenously-tagged allele in *lin-12^Notch^(n676n930)* at 15°C at early L1 stage (left) and early L2 stage (right). **F-G)** Quantification (in %) of Y cell and DA9 neuron in *lin-12^Notch^(n676n930)*, assessed with respectively *ngn-1* reporter from Rashid et al. 2022 (*nsIs913)* and *mig-13* reporters at 15°C (**F**) at early L1 stage (left) and early L2 stage (right), and at 25°C (**G**). Except for 3/92 [1DA9; ngn-1+ “neuronal”] animals (F), all the cells expressing *nsIs913* always also expressed the DA9 marker. **H)** Picture of [1Y; 1DA9] phenotype in WT (top) and [0Y; 2DA9] observed in *lin-12^Notch^(n676n930)* (bottom) early L1 at 15°C assessed with respectively *ngn-1* reporter (*nsIs913*) from Rashid et al. 2022 and *mig-13* reporters. **A-C, E-G)** n, total number of animals scored. Data represent the mean of at least three biological replicates. Two-tailed P value is calculated using a Chi^2^ test. ****P < 0.0001, ***P < 0.001, ***P < 0.01, *P<0.05 **D, H**) Dotted line, rectal slit; white arrowhead, Y cell; red arrowhead, DA9 neurons. The rectal area is pictured. Anterior is to the left and ventral to the bottom.

To re-examine the possibility that we might have overlooked the presence of a “precocious PDA”, we investigated the identity of the Y cell in 15°C *lin-12^Notch^(n676n930)* using a battery of markers. These experiments were conducted using the same protocol and at the precise time point (newly hatched L1) at which Rashid et al.^39^ observed the appearance of a precocious PDA, and we additionally analysed the time of WT Td initiation (23.5h post hatching at 15°C). We first examined two different rectal markers, *egl-5* and *hlh-16,* and never observed a Y cell presenting an axon (purported “precocious PDA”) (Fig6B-C). Instead, at both time points, an absence of the Y cell was observed in most of the worms (Fig6B-D). Since the number of DA9 neurons made in *lin-12^Notch^(n676n930)* was not reported by Rashid et al., we also examined it using the DA9 cell marker *mig-13*^51^. Most *lin-12^Notch^(n676n930)* worms raised at 15°C and 25°C exhibited 2 DA9 neurons, together with a concomitant absence of the Y cell (Fig6C-D), confirming Y^prog^ cell mis-specification into a DA9 neuron. We further used an endogenous *ngn-1* reporter, where GFP has been inserted in-frame at the *ngn-1* endogenous locus. We found *ngn-1* to be expressed only in the WT Y cell, precisely at the time of Td initiation. We thus used it to examine the occurrence of earlier Td initiation in *lin-12^Notch^(n676n930)* mutant raised at 15°C. We never observed any endogenous *ngn-1* expression in the Y cell in newly hatched L1s, either in *lin-12^Notch^(n676n930)* mutant or in WT worms (Fig6E). Later, at the time of Td initiation, only a small proportion of mutant exhibit a Y cell positive for *ngn-1*, confirming that most animals have no Y cell (Fig 6E). Furthermore, *ngn-1* timing of expression in *lin-12^Notch^(n676n930)* mutant occurs just before Td initiation as in WT, in the few animals where a Y cell is made (Fig 6E). Next, we investigated whether the expression of PDA markers starts earlier using the canonical early PDA marker, *cog-1*^28,31–33^. In newly hatched L1 *lin-12^Notch^(n676n930)* mutant grown at 15°C, we never observed a Y cell expressing *cog-1* (SI table1). Finally, since this putative “precocious PDA” had been observed only in the specific *lin-12^Notch^(n676n930)* allele, we examined other *lin-12^Notch^ gf* arrays or alleles using several Y and PDA markers (*egl-5, hlh-16* and *cog-1*). We further confirmed that Notch *gf* activity does not influence the temporality of Y-to-PDA Td (SI table1). Thus, in early L1s, the Y cell never exhibited an axon or expressed a PDA reporter in any of the genetic backgrounds analysed.

We therefore wondered if the precocious neuron observed by Rashid et al. in *lin-12^Notch^(n676n930)* mutant could instead represent mis-specification of the Y^prog^ cell into a DA9 neuron. To confirm this hypothesis, we further looked at the *ngn-1 nsIs913* transgene used in Rashid et al.^39^ whilst also examining a DA9 marker (*mig-13*). In newly hatched *lin-12^Notch^(n676n930)* L1 worms, we observed the presence of an *ngn-1* positive cell with an axon as described by Rashid et al. However, this cell consistently co-expressed the DA9 cell marker, both at 15°C and 25°C (Fig6F-H). In WT worms, we found that the expression profile and dynamics of *nsIs913* is more promiscuous than the endogenously tagged *ngn-1(dev137)* allele. In particular, the endogenous DA9 cell expressed the *ngn-1 nsIs913* reporter at low levels, explaining why the supernumerary DA9 (coming from Y^prog^ cell conversion) also expresses it in *lin-12^Notch^(n676n930)* (Fig6F-H). Confirming these findings, two DA9 neurons were observed at a similar proportion later in the development (Fig6F) when Td initiates, as in newly hatched L1 mutants, and none of the two DA9 neurons expressed the *ngn-1* reporter anymore (not shown). We observed a very small number of early L1s (3/92) presenting a cell with an axon at the characteristic Y position, that was both *ngn-1* positive and *mig-13* negative (“ngn-1+ neuronal” in Fig6F), but at the early L2 stage, this cell was not present anymore. We postulate that these rare events represent failed Y^prog^ to DA9 conversions. As i) we and Rashid et al. observed similar proportions of defective Y cells in *lin-12^Notch^(n676n930)* mutant at 15°C (∼80%, Fig6B-C, Rashid et al^39^); ii) we found that the large majority of *ngn-1* positive cells with an axon are also positive for the DA9 marker; iii) these cells never express a PDA marker; and iv) these cells are not labelled with other Y cells markers, we concluded that the purported “precocious PDA” observed by Rashid et al. is a Y^prog^ cell converted to a DA9 cell. Overall, these results show that the level of Notch signalling does not regulate the timing of Y-to-PDA Td, but rather the cell fate decision between a DA9 neuron and a Y rectal cell during embryogenesis, in parallel to its role in Td.

Altogether, our data suggest a model where the level of Notch signalling impacts the number of Y cells formed, and its timing affects the ability of the Y cell to eventually undergo Td. In absence of the Notch signal, no rectal Y cell is made (and hence 0 PDA neuron is made). WT levels of Notch signalling have two consequences in Y^prog^; it promotes the expression of one set of genes that are necessary to specify Y^prog^ as a rectal cell; and it leads to activation of another subset of genes, including at least *sem-4* and *hlh-16,* which are not involved in making a Y rectal cell at all, but are necessary to endow it with the competence to undergo Td later on. High levels of the Notch signal at the time of Y^prog^ birth result in the formation of 2 Y cells that are both competent and later transdifferentiate into 2 PDA neurons. Finally, high levels of the Notch signal closer to Td initiation result in an absence of PDA formation and the Y cell remains rectal (Fig7).

**Figure 7:**
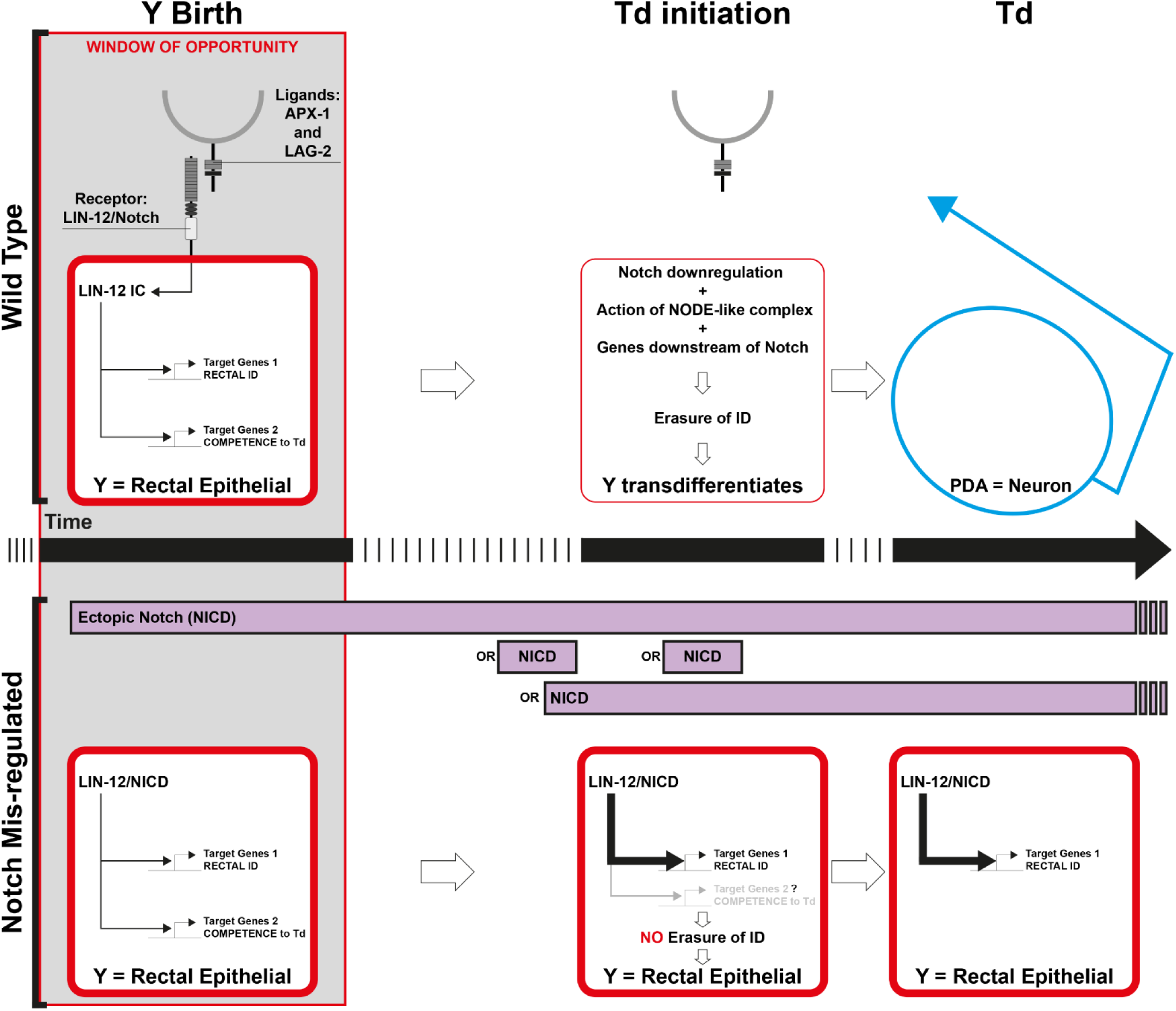
Model for the impact of Notch activity on natural Y-to-PDA. Top half: In the WT situation, a pulse of Notch triggered by two redundant ligands (*apx-1* and *lag-2*) around Y birth will activate two potential and independent sets of genes, endowing the Y cell with its rectal identity as well as its competence to transdifferentiate. The action of these 2 sets can be distinguished, as 2 genes (*sem-4* and *hlh-16*) downstream of Notch are crucial for the initiation of Td while not impacting on Y identity. This time interval of Notch signal represents the window of opportunity (grey box) during which the Notch pathway can promote Td, and a permissive context is required to interpret this Notch signal. In WT animals, Notch signal is then down-regulated at the transcriptional level to allow the Y cell to initiate Td, the first step being the erasure of its initial identity. Here, both the extrinsic environment, as well as the intrinsic cellular context, combine to allow one cell to switch identity. Bottom half: Out-of-time Notch signal after this window of opportunity, either as a continuous signal or as timed pulses (purple rectangles) blocks Y Td. Such late Notch signal does not reset a timer and delay Td, but results in a block of Td by over-imposing in Y a rectal fate. Thus, one signalling pathway, Notch, can exert a dichotomic action, if activated at different steps, on a Td process that occurs in absence of cell division.

## Discussion

Here, we elucidated a novel role of Notch signalling in promoting the competence to transdifferentiate in a natural *in vivo* Td event. Our findings extend a role for Notch in priming cells for Td, beyond induced settings and across species. Indeed, Notch activity has been previously reported to promote experimentally induced reprogramming of germ cells^52^, and also to facilitate the ectopic expression of a specific expression programme, such as muscle or endodermis, in the soma after the forced expression of a relevant cell fate determinant, in *C. elegans*^53^. In addition, Notch signalling promotes Td after injury: for example, the Td of ingrown lymphatic vessels into blood vessels during zebrafish heart regeneration^54^, or the Td of murine pulmonary neuroendocrine cells after lung injury regeneration^55^. Thus, it may be that in several different cellular contexts, the Notch signal triggers the expression of a set of genes that facilitates cell reprogramming.

More precisely, our data suggest that Notch acts on two distinct sets of genes that promote independently (i) the rectal identity, and (ii) the rectal-to-neuronal Td (fig7). Accordingly, we find that the SALL4/*sem-4* and OLIG/*hlh-16* genes lie downstream of the Notch signal and strictly belong to the second set. In *sem-4* and *hlh-16* null mutants a *bona fide* rectal Y cell is made^28^, ^this^ ^study^, but this cell is unable to initiate Td, showing that the two Notch actions can be separated. It is possible that *sem-4* is a direct target of Notch signalling, as several consensus RTGGGAA LAG-1 binding sites^57^, the *C. elegans* CSL protein, can be found in the *sem-4* genomic locus (not shown) and as SALL4/*sem-4* has been suggested to be a *lin-12^Notch^* target during vulval cell specification^56^. However, SEM-4 is only visible in the embryonic Y cell well after the LIN-12^Notch^ receptor has disappeared (Fig2I), a result suggesting that *sem-4* is a secondary -indirect - target of Notch signalling. In addition, it is noteworthy that *sem-4* remains expressed in the Y cell during the whole L1 stage, in absence of a Notch signal, suggesting that *lin-12^Notch^* is not required for its maintenance. Several lines of evidence suggest that *hlh-16/Olig* could be a direct target of the Notch signalling: i) its expression onset occurs immediately after *lin-12^Notch^*expression (Fig 1G-H) (but is maintained until the L1 stage) ii) *hlh-16* has been showed by Bertrand et al.^38^ to be a direct target of the Notch effector LAG-1/Su(H)/CBF. Thus, Notch-dependent events may trigger the expression of these initiation-promoting factors, and possibly set up a positive feedback loop and/or an autoregulatory loop to maintain their expression. Moreover, the Notch signal likely triggers the expression of a number of other unidentified factors, such as earlier competence-related genes, which may in turn positively regulate later initiation-promoting factors.

In addition to triggering the expression of competence and initiation-promoting factors, the *lin-12^Notch^* signal could have additional roles to set up an intrinsic environment prone to the erasure of the rectal identity. It could for instance antagonise Polycomb repressive complex 2 (PRC2) activity, as suggested for *glp-1^Notch^* in the promotion of germline fate^52^. However, while we have previously found a role for the histone demethylase *jmjd3.1* in ensuring the robustness of the later re-differentiation phase of Td^33^, we have not detected a major role for the PRC2 complex in promoting Y rectal identity or antagonising its ability to initiate a Y identity change (AA, S Zuryn & SJ unpublished), suggesting that the importance of the Notch-PRC2 regulatory relationship may vary depending on the process.

While Notch can convert another cell into a transdifferentiation-competent Y cell, most cells in the worm were found refractory to Notch activity, including rectal cells that are seemingly identical to the Y cell, in fact so similar at the transcriptomic level that all rectal cells cluster as one^57^. This highlights the importance of the intrinsic cellular context for the efficiency of a given reprogramming cue. Indeed, the cellular context appears to influence Td in induced settings as well, as over-expression of the muscle determinant MyoD is sufficient to induce the myogenic programme in certain cell types such as adipocytes, but not in all^11^. Similarly, a cocktail of three transcription factors can induce the conversion of pancreatic exocrine cells, but not of the neighbouring cells, into insulin-producing exocrine cells^13^. Why does only DA9^prog^ respond to Notch-induced Td? Making a supernumerary PDA implies two requirements: 1) to have the right toolbox to direct re-differentiation to make a PDA neuron. Indeed, the UNC-3 transcription factor is required for terminal differentiation in both DA neurons and PDA^31,58^, suggesting that DA9^prog^, but not other rectal cells, could be equipped with the right terminal differentiation machinery to make a PDA. And 2) to have a cell that is permissive to interpret the Notch signal resulting in a competent Y cell. One hypothesis is that the permissive context to interpret the Notch signal in the DA9^prog^ cell may represent an evolutionary vestige. It has been suggested that DA9^prog^ and Y^prog^ were multipotent in an ancestral nematode species, forming an equivalence group whose fate was resolved following cell-cell interactions, but that this feature was lost through evolution^17^. It might thus be that these cells have lost their multipotency but have remained equipped with part of the inner machinery underlying this multipotency, a machinery now necessary to dedifferentiate (Fig1A) and initiate Td. Consistently, the DA9^prog^ cell appears to endogenously express several Td initiation-promoting factors, such as *sox-2*, *ceh-6*/oct-4^32^ - known to promote pluripotency in mammals^59^ - and *egl-5*/Hox, which are required to convert a supernumerary Y cell into an extra PDA neuron. These factors may thus be part of DA9^prog^ permissive context.

Interestingly, our data reveal that Notch activity can have a dual effect on the destiny of the Y cell (Fig7). Notch signalling has been reported to exert either positive^e.g.54,55,60–62^ or negative^e.g.63–66^ effects on cellular plasticity in different systems. Indeed, Notch has been reported as essential in various lineages for both self-renewal of progenitor cells and differentiation of their descendants^67–70^. However, here we demonstrated a dichotomic effect of the Notch pathway over time on a single - reprogramming - event. Indeed, according to its timing of activation, Notch signalling can either promote Td (when activated during early embryogenesis) or block it (when activated during the Td initiation). Such dichotomic action on cellular identity dictates that the timing and duration of LIN-12^Notch^ activity should be tightly controlled for proper Td. Our data suggest that this control is at the *lin-12^Notch^* transcriptional level.

How can Notch signalling produce two radically different outcomes? The cellular context of Y^prog^ is likely to change after it has received the Notch signal. Thus, the intrinsic permissive context that combines with an extrinsic Notch signal to endow the Y cell with the competence to transdifferentiate may evolve over time as well. In addition, a late Notch signal results in a definitive block, rather than in a reset of the timing of the process and a later Td initiation. It is likely that prolonged Notch activity reinforces and stabilizes its rectal epithelial identity and makes it so stable that it becomes a barrier to reprogramming. Altogether, our data suggest i) a process that needs some time to occur downstream of the Notch signal and ii) the existence of a window of opportunity for Td to occur. Interestingly, Naylor et al.^6^ reporting on the role of Notch in zebrafish kidney development noted a similar dichotomic effect of Notch overtime: while endogenous Notch promoted the Td of renal epithelial cells into Corpuscles of Stannius gland (CS), overexpressing NICD widely and early diminished the size of the CS. Thus, dichotomic outcomes depending on when Notch activity is seen during a reprogramming event *in vivo* might be a conserved theme. It is possible that in both cases, the Notch signal results in the activation of 2 sets of genes with antagonistic activity; a different gene balance at different time points of the cell’s life may favour one set over the other. The time elapsed between the early Notch pulse and the initiation of Td in WT *C. elegans* is long, corresponding roughly to over 1/3^rd^ of the total developmental time. This time may be necessary to resolve the balance between the instruction to be rectal and the ability to erase this first rectal identity, to ultimately achieve the initiation of dedifferentiation. Does the Notch signal contribute more directly to the initiation of Y Td much later in development? We have ruled out that the levels of this transient extrinsic signal are involved in setting the time of Td initiation. The endogenous Notch pulse in the Y cell could either simply empower the Y cell with the ability to change its identity but have no bearing on the actual initiation of Td; act jointly with a later, unidentified, signal to trigger initiation of Td; or could trigger cell-autonomously a long cascade of events that ultimately result in a dedifferentiation. Further work will be needed to determine what lies downstream of Notch activation and discriminate between these hypotheses.

While biological processes, such as cell-fate acquisition, are often seen as linear and progressive events, our findings once more highlight the importance of the exquisite and highly dynamic regulation of signalling over time. Importantly, our data emphasise how a prolonged (or *out-of-time*) Notch signal could lead to the opposite phenotype to WT. These findings have significant implication for the experimental design of studies aimed at deciphering the role of Notch signalling in biological processes. Our results show that the phenotypes obtained using such experimental designs, and hence conclusions on Notch role, can be opposite to its actual endogenous role. Therefore, the temporality and dynamics of the process studied must be carefully assessed and taken into account.

## Methods

### *C. elegans* maintenance, strains and alleles

Standard methods as described in Brenner 1974^71^ were used for worm handling, maintenance and genetic analysis. Experiments were mostly performed on hermaphrodites at 25°C unless otherwise indicated. The wild-type parent for most strains used in this study is the *C. elegans* var. Bristol strain N2. See SI table 2 for a list of all strains used in this study.

### Microscopic observations and cell identification

DIC and epifluorescence observations were performed using a Zeiss Z1 imager microscope using a Hamamatsu Orca-ER camera C4742 or a Leica DM6 B microscope with LAS X software and the Hamamatsu Digital Camera C11440. Worms were mounted on a 2% agarose pad and anesthetized with 50mM of sodium azide. For all images, anterior is to the left and dorsal up. Single focal planes, or maximum projections when the cells of interest did not coexist on the same focal plane, were used. To assess cell identity, the following criteria were used: for PDA identity (WT PDA or “2PDA” phenotypes), DIC speckled nuclear morphology (note that PDA identification only by DIC is difficult), WT final position, PDA marker expression [*cog-1p::GFP*], presence of an axon projecting posteriorly, then antero-dorsally, as found in WT L3 and older worms. DA9 identity: WT final position, DA9 marker expression [*itr-1p::mCherry* or *mig-13p::mCherry*], presence of an axon projecting posteriorly, then antero-dorsally, as found in WT L4 and older worms. ABplpppaaaa (DA9^prog^), ABprpppaaaa (Y^prog^) and Y cells: DIC “fried egg” nuclear morphology, WT position, marker expression. Of note, another nearby embryonic cell very transiently expresses translational (but not transcriptional) *hlh-16* reporter between ∼1.7-fold and 3-fold embryonic stages. Our scorings in *lin-12^Notch^(gf)* backgrounds were performed before this endogenous expression. Expression of *hlh-16* is maintained in the Y cell until the Td initiation (L2 stage), in WT animals. Fluorescent protein expression was used to follow over-expression of NICD, and mosaic expression associated with non-integrated *Ex* arrays was taken advantage of to analyse cellular focus of action. We have assessed Notch ability to confer the competence to transdifferentiate using PDA markers as final output. While we did not observe a wide plasticity across the worm’s cells, we note that other reprogramming events that would lead to the creation of cells other than a PDA neuron, would go unnoticed. Further studies using a battery of terminal neuronal identities, and other non-neuronal identities, will be required to address this point.

### RNAi experiments

RNAi experiments were performed as previously described. Double stranded RNA (dsRNA) was injected in the relevant genetic background combined to the RNAi hypersensitive mutation *rrf-3(pk1426)*. DsRNA was obtained from *in vitro* transcription of PCR fragment corresponding to cDNA or genomic matrices: *egl-27*, *sem-4, egl-5* and *hlh-16* matrices from Ahringer-MRC feeding RNA interference library^72^: all clones used from this library were first sequence-verified; *ceh-6 target 1* and *sox-2 target 2* matrices from Kagias et al., 2012^32^; *sel-12 :* PCR amplification from genomic DNA using the following primers: F-T7promoter – 5’ taatacgactcactatagggATGCCTTCCACAAGGAGACAAC 3’ and R-T7promoter – 5’ taatacgactcactatagggGAGATCGCTCAAGATATAATCGAAAAG 3’. I*n vitro* transcription was performed using the PCR products as templates with T7 RNA polymerase using the mMESSAGE mMACHINE™ T7 Transcription Kit (Invitrogen™). Single stranded RNA was annealed to form dsRNA by gradually lowering the temperature of the sample from 65°C. *In vitro* transcripts were purified on RNeasy columns (Qiagen) and injected into the gonads and pseudocoelom of young adult worms. F1 progeny derived from these adults were scored for the presence of the Y, PDA and DA9 cells.

### Heat-shock experiments

All the strains used for the heat-shock experiments were maintained at 20°C. Fast induction of fluorescent construct was achieved by raising the temperature as described before^33^ and briefly below.

#### Induction of NICDGFP expression around Y birth

To obtain sufficient numbers of animals to score, egg-pulses were performed with minimum 100 gravid transgenic hermaphrodites at 20°C for one hour on plate with food. Mothers were removed and embryos were left on plate for 15 minutes, when most embryos were between 165 min and 225 min of development. Plates with embryos were sealed and immerged in a water bath at 34°C for 30 minutes. The embryos were immediately cooled down in a 20°C water bath and left to grow at 20°C until the L4 stage, when they were scored. Both transgenic and non-transgenic siblings were handled and scored in parallel.

#### Induction of NICDGFP expression at mid-embryogenesis

Egg-pulses were performed with 100 gravid transgenic hermaphrodites at 20°C for one hour on plate with food. Mothers were removed and embryos were left on plate for 3 hours. Plates were further sealed and immerged in a water bath at 34°C for 30 minutes. The embryos were immediately cooled down in a 20°C water bath and left to grow at 20°C until the L4 stage, when they were scored. Both transgenic and non-transgenic siblings were handled and scored in parallel.

#### Induction of NICDGFP expression before transdifferentiation initiation

Eggs obtained from gravid mothers were allowed to hatch overnight in M9 without food. Synchronized L1 were then put on plate with OP50 for 8 hours before heat-shock at 34°C for 30 minutes. Worms were immediately cooled down in a 20°C water bath and left to grow at 20°C until the L4 stage, when they were scored. Both transgenic and non-transgenic siblings were handled and scored in parallel.

### Temperature-shift experiments

#### Temperature-shift experiments of *glp-1(ar202ts) and glp-1(e2141ts)*

Egg-pulses were performed on pre-warmed plates at 25°C for one hour with gravid hermaphrodites grown at 15°C. Mothers were removed and embryos were left at 25°C (the restrictive temperature) until the adult stage, to score for PDA phenotype and to monitor gonads sterility and morphology.

#### Temperature-shift experiments of *apx-1(zu347ts) and lag-2(q420ts)*

Non-sensitized and sensitized backgrounds were treated the same way during temperature-shift experiments. Worm populations grown at 15°C were bleached. The resulting embryos were put on pre-warmed plates at 26.5°C without food. After 7 hours, all the synchronized hatched L1 larvae were washed away with M9 and discarded, to remove animals that would have been shifted to the restricted temperature too late. OP50 was added subsequently to the plates to allow growth until the L4 stage, when they were scored. Note that the double mutant *lag-2(q420ts) apx-1(zu347ts*)*V* could not be built as the two mutant alleles are genetically very close (0.18 cM).

#### Temperature-shift experiments of *lin-12^Notch^(n676n930)*

Non-sensitized and sensitized backgrounds were treated the same way during temperature-shift experiments. Worms were placed at 15°C or 25°C tightly controlled incubators two generations before scoring. Gravid mothers were bleached and put in suspension in M9 over-night at 15°C or 25°C and place on plates with food the next morning. Worms were directly scored for early L1 scoring, or let at 15°C 23.5h post L1 for early L2 scoring, as described in Rashid et al., 2022^39^.

### SNP mapping of *hlh-16(fp12)*

After the cross between Hawaiian strain WT males and N2 strain *fp12* hermaphrodites, 20 F2 worms with “0 PDA” has been isolated (recombinants *fp12/fp12* homozygotes). Specific SNPs to Hawaiian and N2 strains on chromosome I (positions I-19, I-12, I-6, I5, I14 and I26 cM) have been analysed by PCR/restriction in every 20 recombinants.

### Transgene generation

*C. elegans* extrachromosomal transgenic strains were created by DNA microinjection in the gonad of young adults^73^ of the plasmid of interest together with a co-injection marker and pBSKII^+^ to a final concentration of DNA of 200 ng/µL in water. See SI tables 3 and 4 for a list of all extrachromosomal arrays and integrated arrays, respectively, generated in this study. *hlh-16(syb683[GFP::linker::hlh-16]* (SunyBiotech) was designed by Christelle Gally.

### Plasmid generation

See SI table 5 for a list of all oligonucleotides used in this study.

#### pSJ3177 – MCS::NICDGFP::unc-54 3’UTR

*NICDGFP* was amplified by PCR from the plasmid pSJ201 – *NICDGFP in pBSKII^+^* with the following primers:

F-<u>KpnI</u> – 5’ AAC<u>GGTACC</u>AGAAAAAATGGTTGTTCTGATGTTAGGAGCATTACC 3’

R-<u>KpnI</u> – 5’ TT<u>GGTACC</u>TCAAAAATAATGAGCTGGTTCGGAGTATCG 3’

The obtained PCR product was digested with KpnI and inserted in the multiple cloning site (MCS) 2 of pSJ901 *– MCS1::MCS2::unc-54 3’UTR.*

#### pSJ3171 - hsp-16.2::NICDGFP::unc-54 3′UTR

*hsp-16.2* was excised from pPD49.78 with BamHI and HindIII, and was inserted in MCS1 of pSJ3177 – *MCS::NICDGFP::unc-54 3’UTR.*

#### pSJ6003 - egl-5(6,2kb)Δpes10p::NICDGFP::SL2::mCherry::unc-54 3′UTR

NICDGFP was amplified by PCR from the plasmid pSJ201 with the following primers:

F-<u>KpnI</u> – 5’ AAC<u>GGTACC</u>AGAAAAAATGGTTGTTCTGATGTTAGGAGCATTACC 3’

R-<u>KpnI</u> – 5’ TT<u>GGTACC</u>TCAAAAATAATGAGCTGGTTCGGAGTATCG 3’

The obtained PCR product was inserted in the MCS of pSJ671 - *egl-5(6,7kb)Δpes10p::MCS::SL2::mCherry* at a unique KpnI site.

#### pSJ3173 – col-34p::NICDGFP::unc-54 3′UTR

*col-34p* was amplified by PCR from genomic DNA with the following primers:

F-<u>SphI</u> – 5’ ACA<u>GCATGC</u>GACATGTAAAGTACATCCGTTACATC 3’

R-<u>XmaI</u> – 5’ CCC<u>CCCGGG</u>TGTATGCAGTGGTGGTTTGG 3’

The obtained PCR product was digested by SphI and XmaI, and inserted in the MCS of pSJ3177 – *MCS::NICDGFP::unc-54 3’UTR.*

#### pSJ3169 – lin-48p::NICDGFP::SL2::mCherry::unc-54 3′UTR

*lin-48p was amplified by PCR from genomic DNA with the following primers:*

*F-SphI – 5’* ACAT<u>GCATGC</u>GGATCCAAAAAACCTGCATTTTTTTCAG 3’

R-<u>XmaI</u> – 5’ CCC<u>CCCGGG</u>CTGAAATTGAGCAGAGCTGAAAATTTTTG 3’

The obtained PCR product was digested with SphI and XmaI, and inserted in the MCS of pSJ3177 – *MCS::NICDGFP::unc-54 3’UTR* resulting in pSJ3169 *– lin-48p::NICDGFP::unc-54 3’UTR*

The SL2::mCherry sequence was amplified by PCR from pJG7-psm-SL2-Mcherry (Gift from Cory Bargmann laboratory) with the following primers:

F-<u>NotI</u> – 5’ <u>GCGGCCGC</u>GCTGTCTCATCCTACTTTCACC 3’

R-NotI – 5’ <u>GCGGCCGC</u>CTACTTATACAATTCATCCATGCC 3’

The obtained PCR product was digested by NotI and inserted after NICDGFP in *pSJ3169 – lin-48p::NICDGFP::unc-54 3’UTR.*

#### pSJ3162 – egl-20p::NICDGFP::SL2::mCherry::unc-54 3′UTR

*egl-20p* was amplified by PCR from genomic DNA with the following primers :

F-<u>SphI</u> – 5’ AAA<u>GCATGC</u>GAAGTCATCCTACTAACTAACAATATGACGC 3’

R-<u>XmaI</u> – 5’ AAA<u>CCCGGG</u>TATTTCTGAAATTGAGATGTTTTAGAATTTC 3’

The obtained PCR product was digested with SphI and XmaI, and inserted in the MCS of pSJ3177 – *MCS::NICDGFP::unc-54 3’UTR* resulting in pSJ3164 *– egl-20p::NICDGFP::unc-54 3′UTR*

SL2::mCherry sequenced as been inserted as previous i*n pSJ3164 – egl-20p::NICDGFP::unc-54 3′UTR.*

#### pSJ3103 – egl-5(6,2kb)Δpes10p::NICD::SL2::mCherry::unc-54 3′UTR

The GFP of the NICDGFP sequence was deleted by inverse PCR on pSJ6003 - *egl-5(6,2kb)Δpes10p::NICDGFP::SL2::mCherry::unc-54 3′UTR* with the following primers:

F – 5’ GACTCAACTCATCTGACACCTCC 3’

R – 5’ GGGTCGAGTTACTTTTCTTGAAGG 3’

#### pSJ3215 – col-34p::lin-12cDNA::unc-54 3′UTR and pSJ3217 - col-34p::lin-12cDNA::SL2::mCherry::unc-54 3′UTR

*lin-12* cDNA was amplified from pLM2.4 – *lin-12cDNA in pBSKII^+^* (a gift from the Greenwald laboratory) with the following primers:

F-<u>KpnI</u> – 5’ AAAA<u>GGTACC</u>ATGCGGATCCCTACGATTTG 3’

R-<u>KpnI</u> – 5’ TTTT<u>GGTACC</u>TCAAAAATAATGAGCTGGTTCGG 3’

NICDGFP was excised by KpnI from pSJ3173 – *col-34p::NICDGFP::unc-54 3′UTR* and pSJ3148 - *col-34p::NICDGFP::SL2::mCherry::unc-54 3′UTR* and replaced by the obtained PCR product of *lin-12* cDNA digested by KpnI.

#### pSJ3218 – col-34p::lin-12cDNA(n137)::unc-54 3′UTR

T<u>C</u>T to T<u>T</u>T point mutation was generated by site directed mutagenesis on pSJ3215 - *col-34p::lin-12cDNA::unc-54 3′UTR* with the following primers:

*F-n137 – 5’* GTGTTGTTGACTCAATA<u>TTT</u>GCAAGGCTTGC 3’

R-<u>n137</u> – 5’ GCAAGCCTTGC<u>AAA</u>TATTGAGTCAACAACAC 3’

#### pSJ3222 – col-34p::lin-12cDNA(n941)::unc-54 3′UTR

T<u>G</u>G to T<u>A</u>G Premature stop codon was generated by site directed mutagenesis on pSJ3215 - *col-34p::lin-12cDNA::unc-54 3′UTR* with the following primers:

F-<u>n941</u> – 5’ GGATTCGGTGGGAAA<u>TAG</u>TGTGACGAGCCATTG 3’

R-<u>n941</u> – 5’ CAATGGCTCGTCACA<u>CTA</u>TTTCCCACCGAATCC 3’

#### pSJ3223 – col-34p::lin-12cDNA(ΔANK)::unc-54 3′UTR

Deletion of the seven ankyrin repeats contained in the intracellular part of the *lin-12* receptor was performed by inverse PCR according to the recommendations of Rhett Kovall with the following primers:

F – 5’ CCAGAACGAGAATATTCAATGGATC 3’

R – 5’ AGGTTCAGGTTCAGTTGGAATTTG 3’

#### pSJ3212 – lin-12p::NICDGFP::lin-12UTR

A megaprimer containing NICDGFP was amplified from pSJ3177 with the following primers:

F-5’ CTCAACAGACTTTGCTCAATTTCAAAAAATGGTTGTTCTGATGTTAGGAGCATTAC 3’

R-5’ GGAATTTAAATAATAAATGACGATTGTTCAGAAGATGTACCGAGCTCGGATCCACTAGTAAC 3’

The obtained megaprimer was used to replace the GFP from the plasmid pSJ3143 - *lin-12p::GFPpest-lin-12 1st intron::U54UTR* by Overlap extension PCR cloning.

#### pSJ3240 – lin-12p(ΔR1)::NICDGFP::lin-12 UTR

The first conservation region of the *lin-12* promoter was deleted by inverse PCR on pSJ3212 – *lin-12p::NICDGFP::lin-12UTR* using the following primers:

F-5’ ACAGTAACAGACACCTGTGCTCC 3’

R-5’ TATTGTTAATAAATGAGTGTAACATTTAAG 3’

#### pSJ3242 – lin-12p(ΔR2)::NICDGFPmut::lin-12 UTR

The second conservation region of the *lin-12* promoter was deleted by inverse PCR on pSJ3212 – *lin-12p::NICDGFP::lin-12UTR* using the following primers:

F-5’ ATTAATGATAATGCAAAAGCTACCAGG 3’

R-5’ TGTGTCAGTTTTAGAGTTTTATTTCTG 3’

#### pSJ821 - hlh-16p::mcherry::hlh-16::hlh-16 3’UTR

Translational *hlh-16* reporter from Bertrand et al., 2011^38^ (a gift of Vincent Bertrand/Oliver Hobert) in which GFP was swapped for mCherry.

#### pSJ823 - col-34p::mcherry::hlh-16::hlh-16 3’UTR

*hlh-16* promoter from pSJ821 was replaced by *col-34* promoter.

#### pSJ6334 - ceh-6p::GFP [Transcriptional reporter]

A 5.6kb *ceh-6* promoter fragment was cloned in front of GFP using the following primers :

F-5’ ATAAGAATgcggccgcCGTGTTGCTTTAGCACTTCTCCATCCCTTC

R-5’ ATAGTTTAgcggccgcCAGTTGGGAAGTCCAGGAGCAACGGGGTG

A 3.6kb *ceh-6* 3’UTR fragment was cloned after GFP sop codon using primers :

F-5’ TTTTTTGTGATGCGTATTGATGTAGC

R-5’ GTCGACACAGAAACTACGCAAAATC

#### pSJ1007 - egl-5p(6kb)::2xNLS::mCherry::unc-54 3’UTR 2NLS was added to pSJ671 with the following primers

F- 5’ CGAGCTCAGAAAAAATGACAGCACCGAAAAAAAAGCGAAAAGTTCCAGCTGAG AAGATGACCGCTCCAAAGAAGAAACGCAAAGTA 3’

R- 5’ CCGGTACTTTGCGTTTCTTCTTTGGAGCGGTCATCTTCTCAGCTGGAACTTTTCGCT TTTTTTTCGGTGCTGTCATTTTTTCTGAGCTCGGTAC 3’

#### pSJ834 - egl-5(1,3kb)delta pes10::mkate::unc-54 3’UTR

mkate was added to pSJ6273 using the following primers:

F- 5’ TAGAGGATCCccgGGGATTGGCC

R- 5’ GATATTATACATATTTCATAAAGCCAACC

#### pSJ503- exp-1p::mcherry

An ApaI-BamHI fragment from PD95.75 and containing *mCherry::unc-54 3’UTR* was cloned into the GM4 vector (*exp-1* promotor).

### Statistical analysis

Statistic test was always performed between wild type and mutant. The stars summarize the statistical significance as calculated through the Chi^2^ test. As the Chi^2^ test does not take into account the replicates, no SD or SEM are showed on the graphical representations. However, the data for each experiments are represented by dots on the bar graphs. * p<0.05; ** p<0.01; *** p<0.001; **** p<0.0001 and ns, not significant, for all the tests.

## Acknowledgments

We thank James Priess, Geraldine Maro & Keng Shen, Xiao Liu, Shai Shaham, Stuart Kim and Vincent Bertrand for reagents, Christelle Gally for designing *hlh-16::GFP* CRISPR insertion, Steven Zuryn & Thomas Kleiber for help with experiments and Richard Poole, Peter Meister, Christelle Gally, Thomas Le Gal, Eria Becker and Deborah Warrington for comments on the manuscript. This work was supported by a Ministère de l’Enseignement Supérieur et de la Recherche (to T.D. and J.C.) and an Association pour la Recherche sur le Cancer predoctoral fellowships (to T.D.); and grants from the Fondation pour la Recherche Médicale, the Agence Nationale pour la Recherche (CELLSwitch), the Association Française contre les Myopathies and the Association pour la Recherche sur le Cancer (to S.J.). S.J. is a research director of the CNRS.

## Author contributions

T.D.: Conceptualization, Methodology, Investigation, Formal analysis, Original Draft, Writing - Review & Editing, Visualization ; J.C.: Conceptualization, Methodology, Investigation, Validation, Formal analysis, Original Draft, Writing - Review & Editing, Visualization ; MCM: Methodology, Investigation, Formal analysis; A.A.: Methodology, Investigation, Formal analysis; D.I.: Methodology, Investigation, Formal analysis ; S.J.: Conceptualization, Investigation, Formal analysis, Resources, Writing - Original Draft, Writing - Review & Editing, Supervision, Project administration, Funding acquisition.

