## Supplementary material for "Essential and dual effects of Notch activity on a natural transdifferentiation event": SI figures

| A | Genotype | [1PDA] | [0PDA] | [2PDA] | Sterility | n= |
|---|---|---|---|---|---|---|
| *loss-of-function* | *glp-1(e2141ts)* | 100% | 0% | 0% | 100% | 386 |
| *gain-of-function* | *glp-1(ar202ts)* | 100% | 0% | 0% | 100% | 370 |

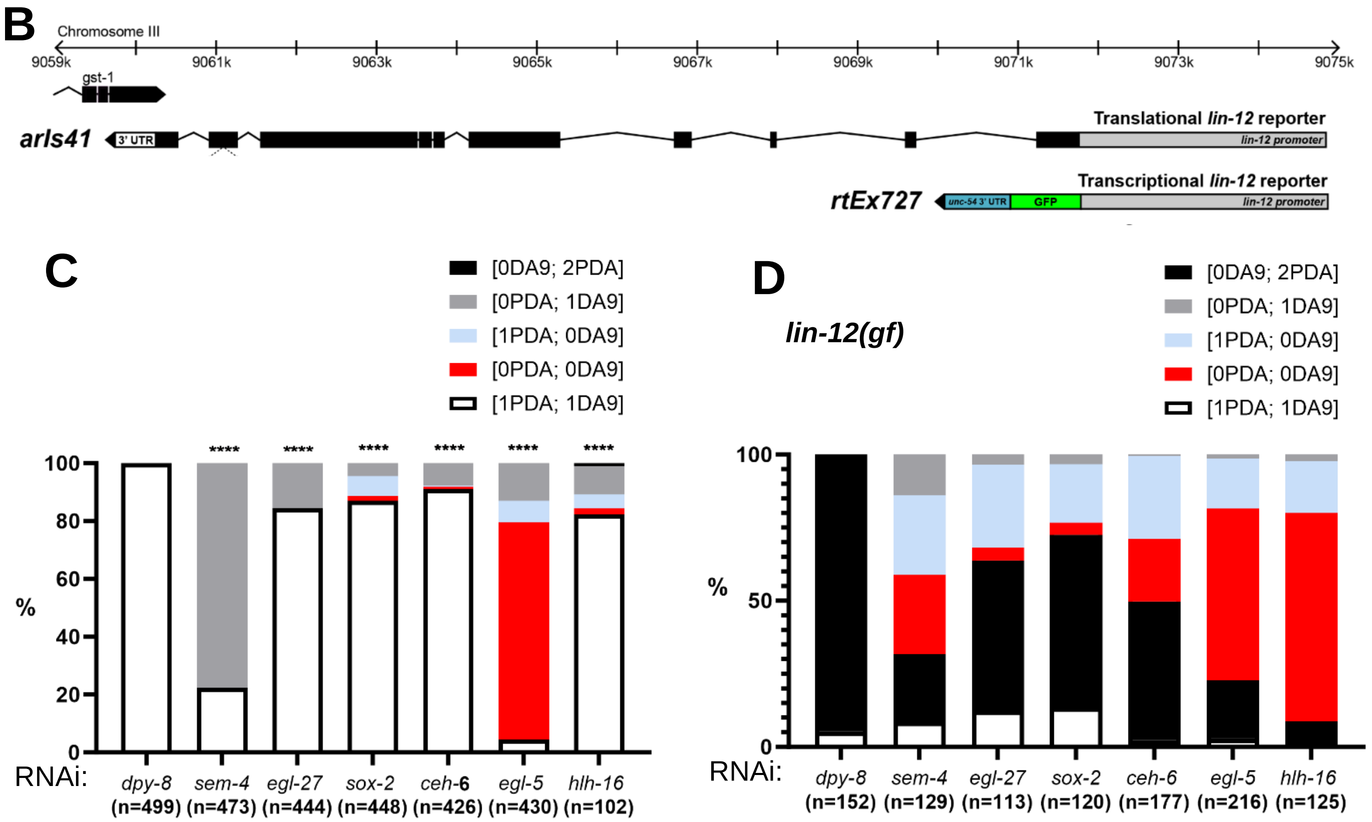


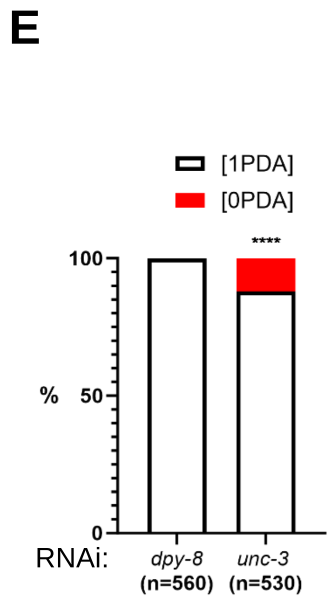


**<u>SI Figure 1:</u>**

A) Y-to-PDA defects (in %) in loss and gain-of-function mutants for *glp-1* at the restrictive temperature (assessed using the *cog-1::gfp* marker). Sterility was assessed to verify that worms have been correctly shifted at the restrictive temperature.

B) Reporters used in Fig1G-H. The translational reporter *arIs41* consists of the entire *lin-12* genomic locus (from 3.5kb upstream of the coding region to 0.8 kb downstream) in which GFP has been inserted in frame in the 9$^{th}$ exon; this translational reporter exhibits rescuing activity (Levitan and Greenwald 1998). The transcriptional reporter *rtEx727*, consists of a 3.5kb promoter fragment of the *lin-12* gene driving the expression of GFP followed by *unc-54* 3'UTR (Singh et al. 2011).

C and D) RNAi knock-down of *dpy-8* (negative control *sem-4*, *egl-27*, *ceh-6*, *sox-2*, *egl-5* and *hlh-16* in (C) the RNAi hypersensitive mutant *rrf-3(pk1426)* and (D) the double mutant *rrf-3(pk1426); lin-12(n950)*, carrying a PDA (*cog-1::gfp*) and a DA9 (*itr-1p::mCherry*) markers.

E) RNAi knock-down of *unc-3* and *dpy-8* (negative control) in the RNAi hypersensitive mutant *rrf-3(pk1426)* carrying a PDA marker (*cog-1::gfp*). Of note, *unc-3*(RNAi) is poorly efficient and leads to a very low penetrance defect.

A, C-E) n, total number of animals scored. The % of each phenotype observed, indicated under brackets above, is represented.

Data in % represent the mean of at least three biological replicates. Two-tailed P value is calculated using a Chi$^2$ test. ****P < 0.0001, ***P < 0.001, ***P < 0.01, *P<0.05.

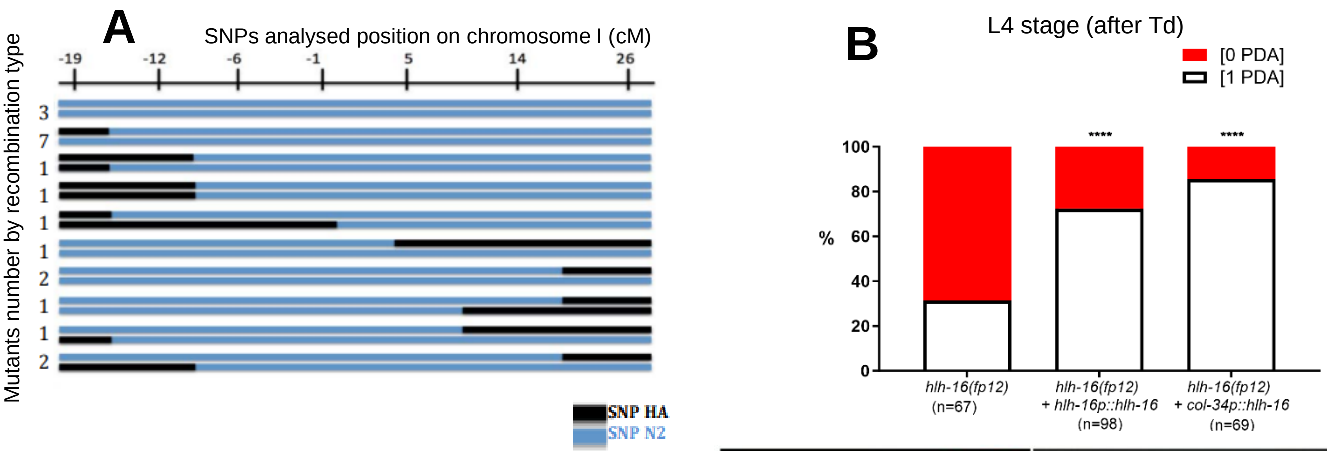


**C** L1 stage (before Td)

| | col-34 | | ceh-6 | |
|---|---|---|---|---|
| | Y cell present | Y cell absent | Y cell present | Y cell absent |
| **WT** | 46 | 0 | 90 | 0 |
| ***hlh-16(fp12)*** | 155 | 0 | 90 | 0 |

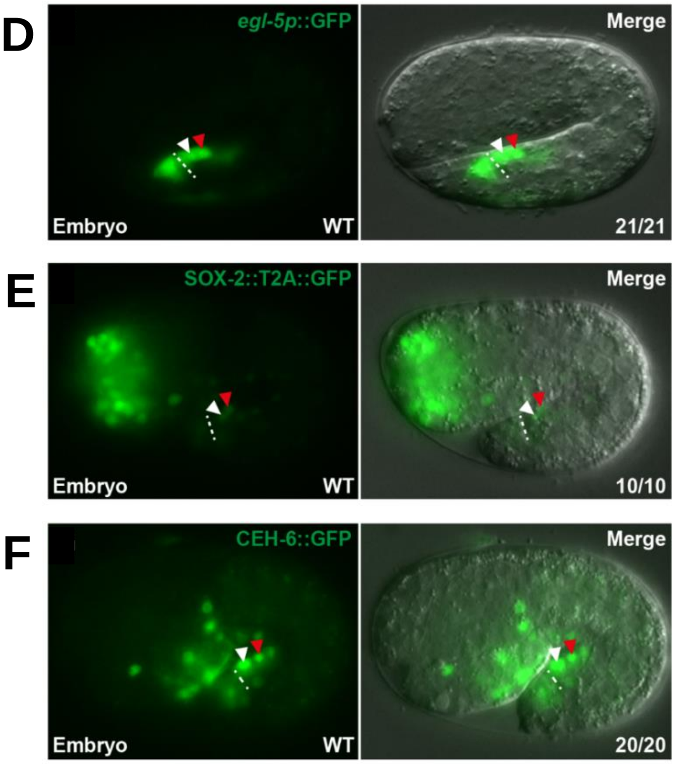


**SI figure 2:**

A) SNPs cartography of the genomic region containing *fp12* allele on chromosome I. SNPs repartition of every recombinant specific to Hawaiian strain (black) and N2 strain (blue) on chromosome I. The number of recombinant mutants is indicated on the left. *fp12* mutation is located between -1cM and 5cM. This region includes the *hlh-16* gene.

B) Rescue of [0 PDA] phenotype observed in *hlh-16(fp12)* loss-of-function mutant by over-expression of *hlh-16* genomic region or over-expression of *hlh-16* under the control of *col-34* (which drives expression in the rectal cells from late embryogenesis (3-fold stage) to Td initiation (Kagias et al. 2012)). *cog-1::gfp*, PDA marker. n, total number of animals scored. Data represent the mean of at least three biological replicates. Two-tailed P value is calculated using a Chi$^2$ test. ****P < 0.0001, ***P < 0.001, ***P < 0.01, *P<0.05

C) Y cell presence at the L1 stage (before the Td initiation) assessed two Y cell markers: *col-34* and *ceh-6*. In both wild type and *hlh-16(fp12)*, the Y rectal cell is always made.

D) *egl-5p*::GFP is expressed in both Y$^{prog}$ and DA9$^{prog}$ cells in WT embryo (1.7-fold).

E) GFP::T2A::SOX-2 fosmid is expressed in both Y$^{prog}$ and DA9$^{prog}$ cells in WT embryo (1.5-fold).

F) CEH-6::GFP fosmid is expressed in both Y$^{prog}$ and DA9$^{prog}$ cells in WT embryo (1.5-fold).

D-F) White arrowhead: Y cell; red arrowhead: DA9. Dotted line: rectal slit. Numbers indicates the fraction of worms displaying this representative expression pattern over the total number of worms scored. Anterior =left. Left panel = fluorescence microscopy; right panel = merged fluorescence and DIC microscopy.

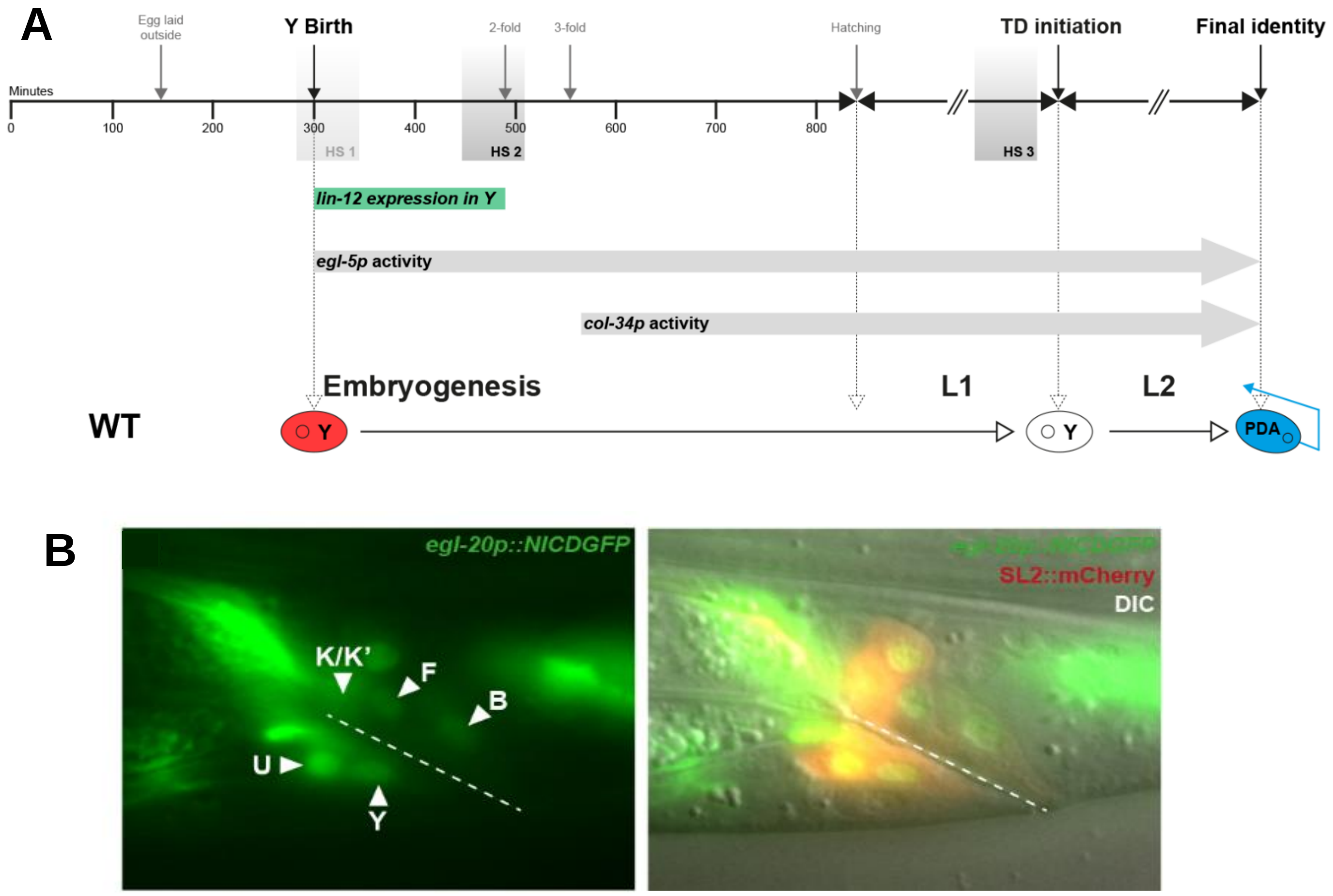


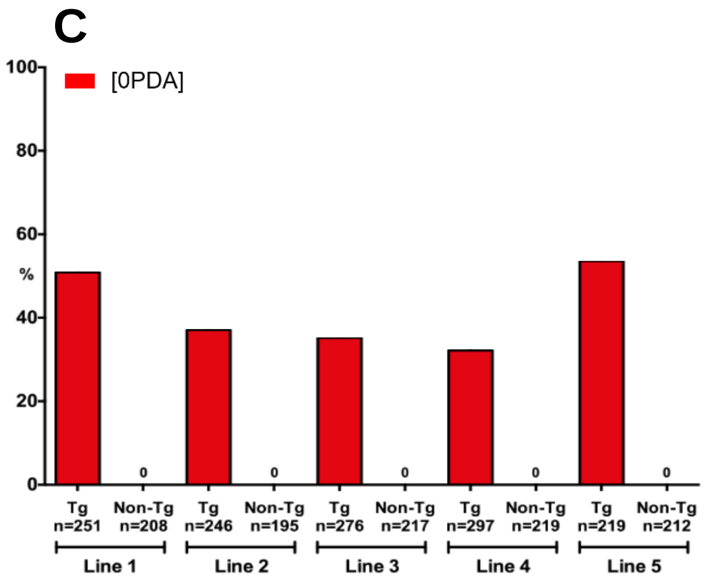


**SI Figure 3:**

**A)** Timing of expression of the *lin-12*, *egl-5* and *col-34* genes or drivers used, and of the heat-shock (HS) 1, 2 and 3 with respect to the timeline of the different steps of the Y-to-PDA Td.

**B)** Wide-spread mosaic expression of *egl-20p*::*NICDGFP* in all the rectal cells including in the Y cell. Representative pattern in a L4 transgenic worm. Anterior = left ; ventral = bottom.

**C)** Quantification (in %) of [0 PDA] (Td defect) in five independent transgenic lines expressing *egl-20p::NICDGFP::SL2mCherry*.

Tg, transgenic worms; Non-tg, non-transgenic siblings. n, total number of animals scored. Data represent the mean of at least three biological replicates. Two-tailed P value is calculated using a Chi$^2$ test. ****P < 0.0001, ***P < 0.001, ***P < 0.01, *P<0.05

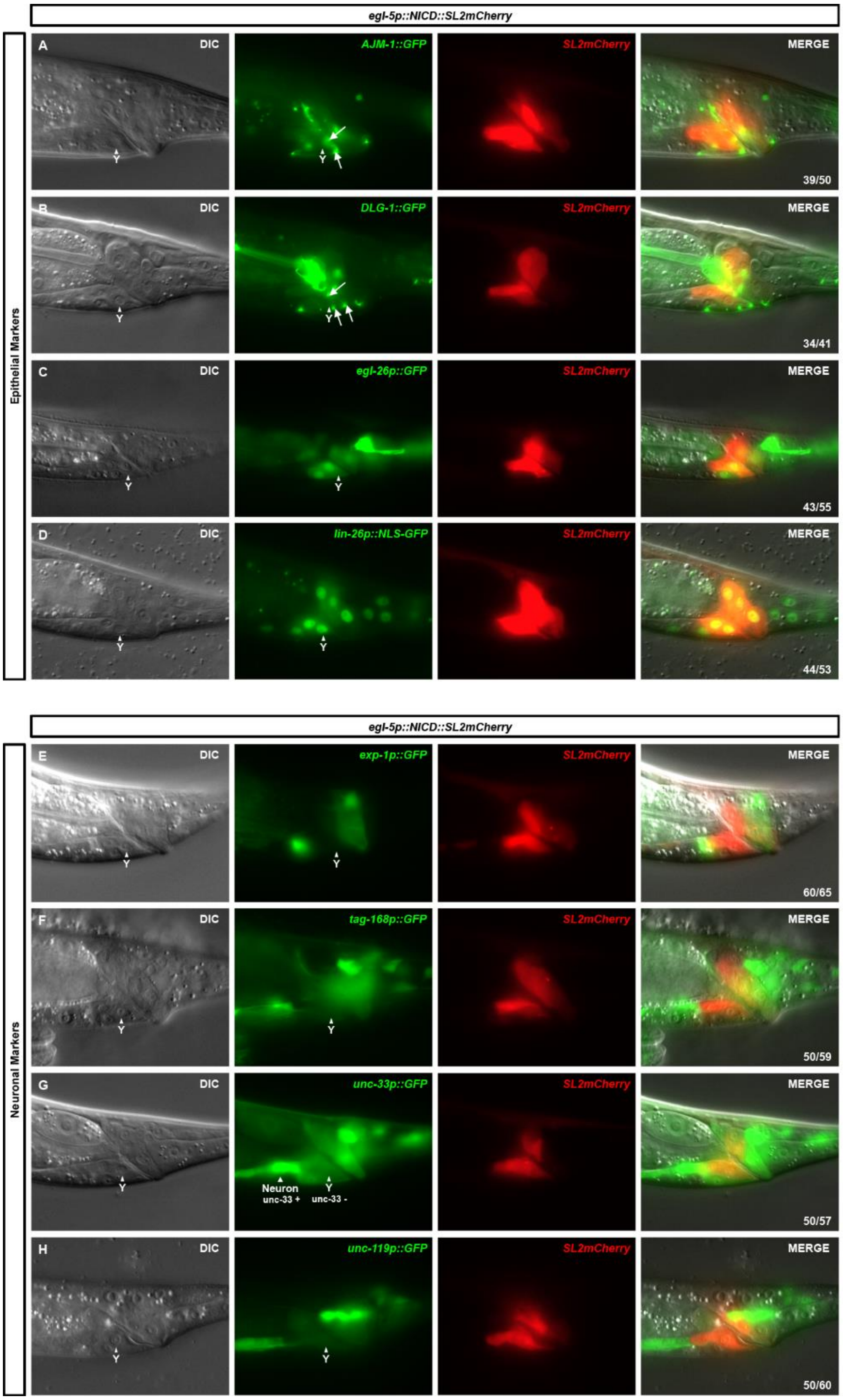


**SI Figure 4:**

Representative pictures of different epithelial (upper panels) or neuronal (lower panels) markers expression in the persistent Y cell when LIN-12$^{Notch}$ signal is maintained throughout (using the *egl-5p::NICD::SL2::mCherry* transgene). DIC (left), epifluorescence (middle) and merged (right) images in L4 transgenic worms are shown. SL2::mCherry shows NICD expression in the rectal cells. Numbers represent the fraction of worms showing this representative phenotype over the total number of animals scored. Anterior is to the left and ventral to the bottom.

A-B) epithelial junction markers, (A) *AJM- 1::GFP* and (B) *DLG- 1::GFP* are expressed (arrows) in the persistent Y (arrowhead).

C) The rectal cell marker, *egl-26p*::*GFP*, is expressed in the persistent Y (arrowhead).

D) The epithelial differentiation marker, *lin-26p::NLS-GFP*, is expressed in the persistent Y (arrowhead).

E) The PDA marker, *exp-1p*::*GFP*, is not expressed in the persistent Y (arrowhead).

F-G) The pan-neuronal marker (F) *tag-168p::GFP* and (G) *unc-33p::GFP* are not expressed in the persistent Y (arrowhead). Note that *unc-33p::GFP* displays a high background around positive cells. Nevertheless, the difference between an *unc-33* positive cell and an *unc-33* negative cell is clear.

H) The pan-neuronal marker *unc-119p::GFP*, is not expressed in the persistent Y (arrowhead).

**A**

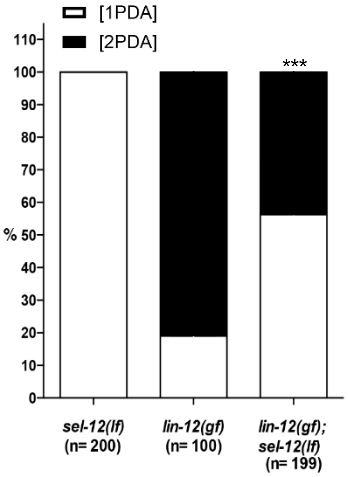


**B**

| | Genotype | RNAi | T°C | PDA OK | No PDA | n = |
|---|---|---|---|---|---|---|
| DSL | ***arg-1***(*ok3127)X* | - | 25°C | 100% | 0% | 582 |
| | *rrf-3(pk1426)II;* ***arg-1***(*ok3127)X* | *sel-12* | 20°C | 100% | 0% | 582 |
| | ***dsl-1***(*ok810)III* | - | 25°C | 100% | 0% | 367 |
| | ***dsl-1***(*ok810)III; sel-12(ar171)unc-1(e538)X* | - | 25°C | 100% | 0% | 441 |
| | ***dsl-3***(*ok3411)IV* | - | 25°C | 100% | 0% | 389 |
| | ***dsl-3***(*ok3411)IV; sel-12(ar171)unc-1(e538)X* | - | 25°C | 100% | 0% | 287 |
| | ***dsl-4***(*ok1020)X* | - | 25°C | 100% | 0% | 285 |
| | *rrf-3(pk1426);* ***dsl-4***(*ok1020)X* | *sel-12* | 20°C | 100% | 0% | 697 |
| | ***dsl-5***(*ok588)IV* | - | 25°C | 100% | 0% | 349 |
| | ***dsl-5***(*ok588)IV; sel-12(ar171)unc-1(e538)X* | - | 25°C | 100% | 0% | 344 |
| | ***dsl-6***(*ok2265)IV* | - | 25°C | 100% | 0% | 310 |
| | ***dsl-6***(*ok2265)IV; sel-12(ar171)unc-1(e538)X* | - | 25°C | 100% | 0% | 252 |
| DOS | ***osm-7***(*n1515)III* | - | 25°C | 100% | 0% | 253 |
| | ***osm-7***(*n1515)III; sel-12(ar171)unc-1(e538)X* | - | 25°C | 100% | 0% | 308 |
| | ***osm-11***(*rt192)X* | - | 25°C | 100% | 0% | 396 |
| | *rrf-3(pk1426)II;* ***osm-11***(*rt192)X* | *sel-12* | 20°C | 100% | 0% | 412 |
| | ***dos-1***(*ok2398)III* | - | 25°C | 100% | 0% | 353 |
| | ***dos-1***(*ok2398)III; sel-12(ar171)unc-1(e538)X* | - | 25°C | 100% | 0% | 299 |
| | ***dos-2***(*tm4515)II* | - | 25°C | 100% | 0% | 480 |
| | ***dos-2***(*tm4515)II; sel-12(ar171)unc-1(e538)X* | - | 25°C | 100% | 0% | 306 |
| | ***dos-3***(*tm4899)III* | - | 25°C | 100% | 0% | 425 |
| | ***dos-3***(*tm4899)III; sel-12(ar171)unc-1(e538)X* | - | 25°C | 100% | 0% | 393 |

**SI Figure 5:**

**A)** Quantification (in %) of [2 PDA] in *sel-12(ar171)* single mutant [noted *sel-12(lf)*] compared to *lin-12(n950)* gain-of-function [noted *lin-12(gf)*] and double *lin-12(n950);sel-12(ar172)* mutants. *sel-12* mutant is WT while a double mutant *lin-12(n950); sel-12(ar171)* exhibits a reduction of [2 PDA] caused by *lin-12(n950),* and has therefore been hereafter used as a sensitised background for Notch signalling deficiencies. Data represent the mean of two biological replicates for *lin-12(gf)* and four biological replicates for *sel-12(lf)* and *lin-12(gf);sel-12(lf)* mutants. Two-tailed P value is calculated using a Chi$^2$ test; ***P < 0.001.

**B)** Scoring of all the available mutants for non-canonical ligands, alone or associated with *sel-12(ar171)* at the indicated temperature, and expressed in % of total animals scored (n). Data represent biological triplicates.

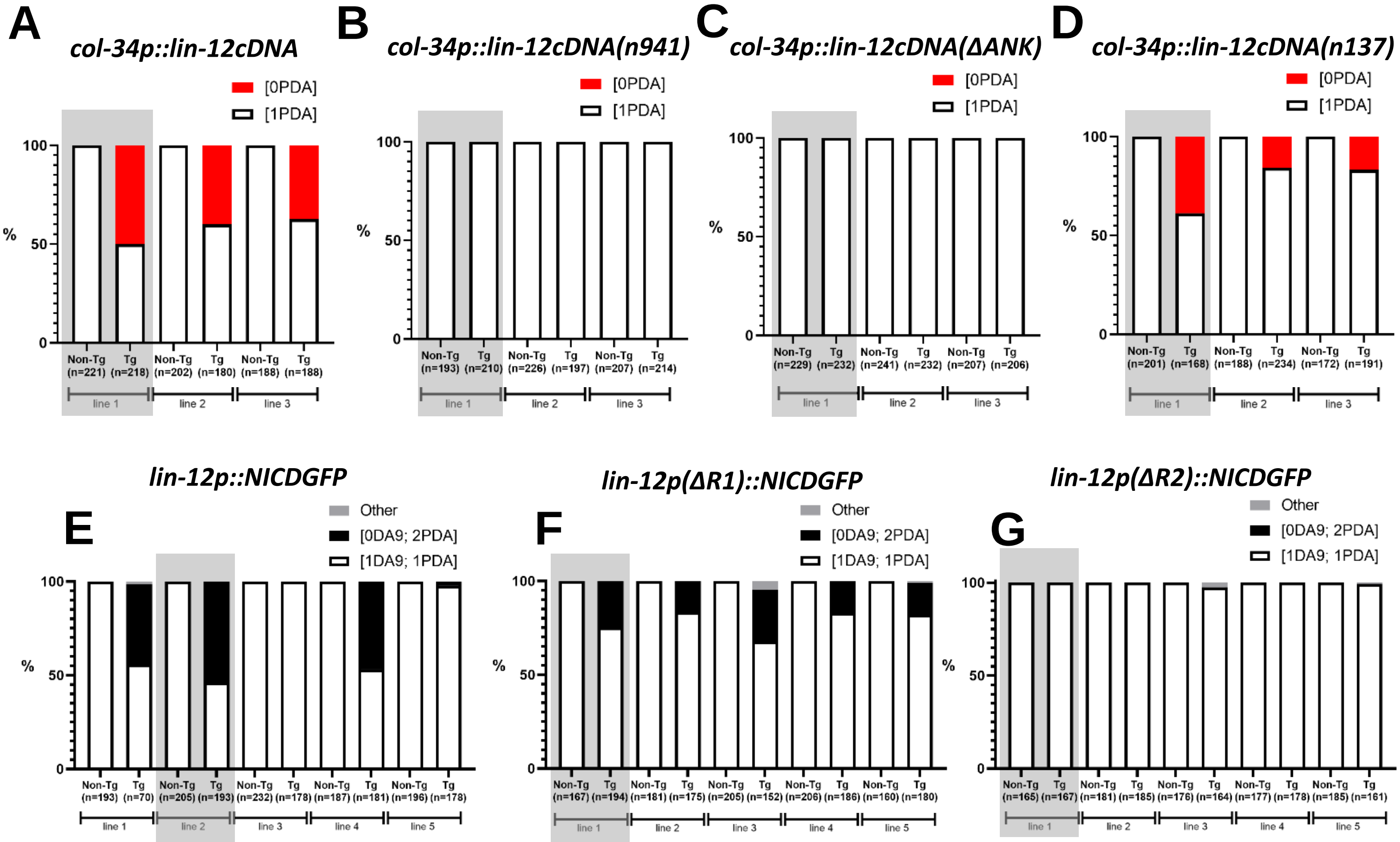


**<u>SI Figure 6:</u>**

**A)** Quantification of the [0 PDA] phenotype in lines carrying a *col-34p*::*lin-12(WT)*cDNA transgene. When expression of *lin-12* is prolonged until the initiation of Td, Y Td is impaired and a [0 PDA] phenotype appears in transgenic (Tg) worms but not in non-transgenic (non-Tg) control siblings.

**B)** Quantification of the No Td [0 PDA] phenotype in lines carrying the negative control transgene *col-34p::lin-12cDNA(n941).* All the scored worms are wild type.

**C)** Quantification of the [0 PDA] phenotype in lines carrying the negative control transgene *col-34p*::*lin-12cDNA(ΔANK)*. All the scored worms are wild type.

**D)** Quantification of the [0 PDA] phenotype in lines carrying the positive control *col-34p::lin-12cDNA(n137).* Transgenic worms display a [0 PDA] phenotype.

**A-D)** Three independent transgenic lines were scored (in %) for each construct. The grey boxes represent the lines depicted in the Figure 5E. Data represent the mean of three biological replicates. Tg, transgenic worms. Non-Tg, non-transgenic control siblings. PDA presence is assessed using *cog-1::gfp.*

**E)** Quantification of the number of PDA and DA9 neurons in lines expressing *lin-12p*::NICD. A supernumerary TD [2 PDA] phenotype appears in transgenics worms in 4 out of 5 lines while all the non-transgenics siblings are WT.

**F)** Quantification of the number of PDA and DA9 neurons in lines expressing *lin-12p*(ΔR1)::NICD. Deletion of the R1 region does not abolish transgene activity, as a [2 PDA] phenotype can still be obtained.

**G)** Quantification of the number of PDA and DA9 neurons in lines expressing *lin-12p*(ΔR2)::NICD. Transgene expression appears affected by deletion of the R2 region, as exemplified by the absence of a [2 PDA] phenotype.

**E-G)** Five independent transgenic lines were scored (in %) for each construct. The grey boxes represent the lines depicted in the Figure 5F. Data represent the mean of three biological replicates. Tg, transgenic worms; Non-Tg, non-transgenic siblings. PDA and DA9 presence are assessed with *cog-1::gfp* and *itr-1p::mCherry* reporters respectively.
