## Supplementary material for "Essential and dual effects of Notch activity on a natural transdifferentiation event": SI tables

**SI Table 1: No « precaucious PDA » is observed in newly hatched L1 with elevated Notch activity, as assessed using *cog-1* expression**

| Markers | *cog-1* [&] | *hlh-16* [#] | *egl-5* [#] |
|---|---|---|---|
| *lin-12(n676n930) 15°C* | 96/96 no expression | - | - |
| *lin-12(n302)* | 106/106 no expression | - | - |
| *lin-12(n950)* | 82/82 no expression | - | - |
| *lin-12p::NICD* | 100/100 no expression | 46/91 (51%) of [2Y] | 46/91 (51%) of [2Y] |

&, Data expressed as n animal with NO expression / total number of animals scored
[#], Data expressed as n animal with 2 Y cells expressing the marker / total number of animals scored

**SI Table 2: *C. elegans* strains used in this study**

Note that the transgenes generated in this study are followed by "*". Refer to SI tables 3 (extrachromosomal arrays) and 4 (integrated arrays) for more details.

| Strain Name | Purpose | Reference |
|---|---|---|
| Wild-type background | | |
| IS98 | *syIs63[cog-1::gfp;unc-119(+)] IV (outcrossed 8x)* | Jarriault *et al.*, 2008 |
| *lin-12* mutant characterisation | | |
| IS791 | *lin-12(n950) III; syIs63[cog-1::gfp;unc-119(+)] IV* | Jarriault *et al.*, 2008 |
| IS164 | *lin-12(n302) III; syIs63[cog-1::gfp;unc-119(+)] IV* | This study |
| IS2120 | *syIs63[cog-1::gfp;unc-119(+)] IV; fpEx683[mig-13p::mCherry; myo-2::mCherry]** | This study |
| IS2121 | *syIs63[cog-1::gfp;unc-119(+)] IV; fpEx684[mig-13p::mCherry; myo-2::mCherry]** | This study |
| IS2123 | *syIs63[cog-1::gfp;unc-119(+)] IV; fpEx686[mig-13p::mCherry; myo-2::mCherry]** | This study |
| IS2118 | *lin-12(n950) III; syIs63[cog-1::gfp;unc-119(+)] IV; fpEx682[mig-13p::mCherry; myo-2::mCherry]** | This study |
| IS2115 | *syIs63[cog-1::gfp;unc-119(+)] IV; wyEx1902[itr-1p::mCherry; odr-1p::GFP]* | This study |
| IS2116 | *lin-12(n950) III; syIs63[cog-1::gfp;unc-119(+)] IV; wyEx1902[itr-1p::mCherry; odr-1p::GFP]* | This study |
| IS2199 | *syIs63[cog-1::gfp;unc-119(+)] IV; fpIs67[itr-1p::mCherry; odr-1p::GFP]** | This study |
| IS2235 | *unc-32(e189) lin-12(n676n930)III; syIs63[cog-1::gfp;unc-119(+)] IV; fpIs67[itr-1p::mCherry; odr-1p::GFP]** | This study |

| | | |
|---|---|---|
| *glp-1* mutant characterisation | | |
| IS790 | *glp-1(e2141ts) III; syIs63[cog-1::gfp;unc-119(+)] IV* | This study |
| IS2153 | *glp-1(ar202ts) IIII; syIs63[cog-1::gfp;unc-119(+)] IV* | This study |
| *hsp::NICDGFP* transgenic strains | | |
| IS2093 | *syIs63[cog-1::gfp;unc-119(+)] IV; fpEx664[hsp-16.2::NICDGFP; myo-2::mCherry]** | This study |
| IS2094 | *syIs63[cog-1::gfp;unc-119(+)] IV; fpEx665[hsp-16.2::NICDGFP; myo-2::mCherry]** | This study |
| IS2095 | *syIs63[cog-1::gfp;unc-119(+)] IV; fpEx666[hsp-16.2::NICDGFP; myo-2::mCherry]** | This study |
| RNAi sensitised background | | |
| IS85 | *rrf-3(pk1426) II; syIs63[cog-1::gfp;unc-119(+)] IV* | Kagias et al., 2012 |
| IS1676 | *rrf-3(pk1426) II ; lin-12(n950) III ; syIs63[cog-1::gfp;unc-119(+)] IV* | This Study |
| IS2274 | *rrf-3(pk1426) II ; fpIs67[odr-1:GFP; itr-1p::mcherry]* ; syIs63[cog-1::gfp;unc-119(+)] IV* | This study |
| IS2286 | *rrf-3(pk1426) II ; lin-12(n950) III ; fpIs67[odr-1:GFP; itr-1p::mcherry]* ; syIs63[cog-1::gfp;unc-119(+)] IV* | This study |
| *hlh-16* mutant characterisation | | |
| IS3 | *bxIs7[egl-5::gfp; lin-15(+)] I* | Jarriault *et al.*, 2008 |
| IS3442 | *hlh-16(fp12) I; bxIs7[egl-5::gfp; lin-15(+)] I* | This study |
| IS663 | *hlh-16(fp12) I; [cog-1::gfp;unc-119(+)] IV* | This Study |
| IS2330 | *fpEx828[hlh-16p::mcherry::hlh-16::hlh-16 3'UTR(20ng); myo-2::GFP(3ng)] * ; hlh16(fp12) I ; syIs63[cog-1::gfp;unc-119(+)] IV* | This study |
| IS2332 | *fpEx830[col-34p::mcherry::hlh-16::hlh-16 3'UTR(20ng); myo-2::GFP(3ng)] * ; hlh16(fp12) I ; syIs63[cog-1::gfp;unc-119(+)] IV* | This study |
| IS1299 | *gaIs245[col-34p::HIS-24::mCherry; unc-119(+)] ; oxIs12[unc-47::gfp; lin-15(+)] X* | Riva et al., 2022 |
| IS3126 | *hlh-16(fp12) I; gaIs245[col-34p::HIS-24::mCherry; unc-119(+)] V; oxIs12[unc-47::gfp; lin-15(+)] X* | This study |
| IS2677 | *hlh-16(fp12) I; fpEx929[ceh-6 locus 13kb (ceh-6 ORF removed= only GFP/transcriptional in strataclone/sens1 -10ng/µl); myo-2::GFP 2ng/µl; pBluescript 200ng/µl]** | This study |
| IS2592 | *fpEx929[ceh-6 locus 13kb [ceh-6 ORF removed= only GFP/transcriptional] in strataclone/sens1 -10ng/µl; myo-2::GFP 2ng/µl; pBluescript 200ng/µl]* | This study |
| IS318 | *hlh-16(fp12) I; syIs63[cog-1::gfp;unc-119(+)] IV* | This study |
| CB4856 | *Hawaiien wild type strain* | WBPaper00005369 |
| *lin-12* reporter strains | | |

| | | |
|---|---|---|
| GS4335 | *arIs41[LIN-12::gfp; pRF4]* | Levitan *et al.*, 1998 |
| HA2182 | *pha-1(e2123)III; rtEx727[lin-12p::gfp, myo-2p::gfp, pha-1(+)]* | Singh *et al.*, 2011 |
| *promoter-specific::NICDGFP* transgenic strains | | |
| *egl-5p::NICDGFP* | | |
| IS834 | *syIs63[cog-1::gfp;unc-119(+)] IV; fpEx84[egl-5(6,2kb)Δpes10p::NICDGFP::SL2::mCherry, myo-2p::GFP]** | This study |
| IS835 | *syIs63[cog-1::gfp;unc-119(+)] IV; fpEx85[egl-5(6,2kb)Δpes10p::NICDGFP::SL2::mCherry, myo-2p::GFP]** | This study |
| IS836 | *syIs63[cog-1::gfp;unc-119(+)] IV; fpEx86[egl-5(6,2kb)Δpes10p::NICDGFP::SL2::mCherry, myo-2p::GFP]** | This study |
| IS837 | *syIs63[cog-1::gfp;unc-119(+)] IV; fpEx87[egl-5(6,2kb)Δpes10p::NICDGFP::SL2::mCherry, myo-2p::GFP]** | This study |
| IS838 | *syIs63[cog-1::gfp;unc-119(+)] IV; fpEx88[egl-5(6,2kb)Δpes10p::NICDGFP::SL2::mCherry, myo-2p::GFP]** | This study |
| *col-34p::NICDGFP* | | |
| IS1052 | *syIs63[cog-1::gfp;unc-119(+)] IV; fpEx217[col-34p::NICDGFP; myo-2p::GFP] ** | This study |
| IS1053 | *syIs63[cog-1::gfp;unc-119(+)] IV; fpEx218[col-34p::NICDGFP; myo-2p::GFP] ** | This study |
| IS1054 | *syIs63[cog-1::gfp;unc-119(+)] IV; fpEx219[col-34p::NICDGFP; myo-2p::GFP] ** | This study |
| IS1055 | *syIs63[cog-1::gfp;unc-119(+)] IV; fpEx220[col-34p::NICDGFP; myo-2p::GFP] ** | This study |
| IS1056 | *syIs63[cog-1::gfp;unc-119(+)] IV; fpEx221[col-34p::NICDGFP; myo-2p::GFP] ** | This study |
| *lin-48p::NICDGFP* | | |
| IS1388 | *syIs63[cog-1::gfp;unc-119(+)] IV; fpEx334[lin-48p::NICDGFP::SL2::mCherry; myo-2p::GFP]** | This study |
| IS1389 | *syIs63[cog-1::gfp;unc-119(+)] IV; fpEx335[lin-48p::NICDGFP::SL2::mCherry; myo-2p::GFP]** | This study |
| IS1390 | *syIs63[cog-1::gfp;unc-119(+)] IV; fpEx336[lin-48p::NICDGFP::SL2::mCherry; myo-2p::GFP]** | This study |
| IS1391 | *syIs63[cog-1::gfp;unc-119(+)] IV; fpEx337[lin-48p::NICDGFP::SL2::mCherry; myo-2p::GFP]** | This study |
| IS1392 | *syIs63[cog-1::gfp;unc-119(+)] IV; fpEx338[lin-48p::NICDGFP::SL2::mCherry; myo-2p::GFP]** | This study |
| *egl-20p::NICDGFP* | | |

| | | |
|---|---|---|
| IS1623 | *syIs63[cog-1::gfp;unc-119(+)] IV; fpEx459[egl-20p::NICDGFP::SL2::mCherry; myo-2p::mCherry]** | This study |
| IS1624 | *syIs63[cog-1::gfp;unc-119(+)] IV; fpEx460[egl-20p::NICDGFP::SL2::mCherry; myo-2p::mCherry] ** | This study |
| IS1625 | *syIs63[cog-1::gfp;unc-119(+)] IV; fpEx461[egl-20p::NICDGFP::SL2::mCherry; myo-2p::mCherry] ** | This study |
| IS1626 | *syIs63[cog-1::gfp;unc-119(+)] IV; fpEx462[egl-20p::NICDGFP::SL2::mCherry; myo-2p::mCherry] ** | This study |
| IS1627 | *syIs63[cog-1::gfp;unc-119(+)] IV; fpEx463[egl-20p::NICDGFP::SL2::mCherry; myo-2p::mCherry] ** | This study |
| | *egl-5p::NICD* | |
| IS1831 | *fpIs51[egl-5(6,2kb)Δpes10p::NICD::SL2::mCherry] ** | This study |
| IS1880 | *syIs63[cog-1::gfp;unc-119(+)] IV; fpIs51[egl-5(6,2kb)Δpes10p::NICD::SL2::mCherry] ** | This study |
| | Reporter strains used to assess Y identity | |
| IS1908 | *fpIs51[egl-5(6,2kb)Δpes10p::NICD::SL2::mCherry]*; mcIs17[lin-26p::NLS-GFP;pRF4]* | This study |
| IS1900 | *fpIs51[egl-5(6,2kb)Δpes10p::NICD::SL2::mCherry]*; kuIs36[egl-26p::gfp; unc-119(+)]* | This study |
| IS1904 | *jcIs1[AJM-1::GFP; pRF4] IV; fpIs51[egl-5(6,2kb)Δpes10p::NICD::SL2::mCherry] ** | This study |
| IS1902 | *fpIs51[egl-5(6,2kb)Δpes10p::NICD::SL2::mCherry] *; mcIs47[DLG-1::gfp; pRF4]* | This study |
| IS1912 | *otIs117[unc-4(+); unc-33p::GFP] IV; fpIs51[egl-5(6,2kb)Δpes10p::NICD::SL2::mCherry]** | This study |
| IS1911 | *ncIs3[tag-168p::GFP] III; fpIs54[egl-5(6,2kb)Δpes10p::NICD::SL2::mCherry] ** | This study |
| IS1910 | *edIs6[unc-119p::gfp; rol-6] IV; fpIs51[egl-5(6,2kb)Δpes10p::NICD::SL2::mCherry] ** | This study |
| IS1915 | *wyIs75[unc-47p::DsRed; exp-1p::GFP; odr-1p::RFP] III; fpIs51[egl-5(6,2kb)Δpes10p::NICD::SL2::mCherry] ** | This study |
| IS1916 | *wyIs75[unc-47p::DsRed; exp-1p::GFP; odr-1p::RFP] III; fpIs54[egl-5(6,2kb)Δpes10p::NICD::SL2::mCherry]** | This study |
| | Reporter strains used to assess rectal cells identity | |
| IS1908 | *fpIs51[egl-5(6,2kb)Δpes10p::NICD::SL2::mCherry] *; mcIs17[lin-26p::NLS-GFP;pRF4]* | This study |
| IS2198 | *fpIs51[egl-5(6,2kb)Δpes10p::NICD::SL2::mCherry] *; saIs14[lin-48p::GFP; unc-119(+)]* | This study |

| | | |
|---|---|---|
| IS2149 | *eIs34[mab-9p::gfp; pCes1943 (rol-6D)] III; fpIs51[egl-5(6,2kb)Δpes10p::NICD::SL2::mCherry] ** | This study |
| Canonical Notch ligand mutant strains | | |
| IS794 | *syIs63[cog-1::gfp;unc-119(+)] IV; sel-12(ar171) unc-1(e538) X* | This study |
| IS792 | *lin-12(n950 III; syIs63[cog-1::gfp;unc-119(+)] IV; sel-12(ar171) unc-1(e538) X* | This study |
| IS1361 | *syIs63[cog-1::gfp;unc-119(+)] IV; apx-1(zu347ts)V* | This study |
| IS1362 | *syIs63[cog-1::gfp;unc-119(+)] IV; apx-1(zu347ts)V; sel-12(ar171) unc-1(e538) X* | This study |
| IS1364 | *syIs63[cog-1::gfp;unc-119(+)] IV; lag-2(q420ts)V* | This study |
| Ligand reporter strains | | |
| GS3795 | *dpy-20(e1282) IV; arIs98[apx-1p::2NLS::YFP; ceh-22::gfp; pMH86]* | Li *et al.*, 2010 |
| JK2049 | *qIs19[lag-2p::GFP; pRF4] V* | Blelloch *et al.*, 1999 |
| Non-canonical ligand mutant strains | | |
| IS1925 | *syIs63[cog-1::gfp;unc-119(+)] IV; arg-1(ok3127)X* | This study |
| IS2009 | *rrf-3(pk1426) II; syIs63[cog-1::gfp;unc-119(+)] IV; arg-1(ok3127)X* | This study |
| IS770 | *dsl-1(ok810) V; fpIs11[exp-1:p:mCherry; myo-2p::GFP]** | This study |
| IS973 | *dsl-1(ok810) IV; sel-12(ar171) unc-1(e538) X; fpIs10[exp-1p::mCherry; myo-2p::GFP]** | This study |
| IS1895 | *wyIs75[unc-47p::DsRed; exp-1p::GFP; odr-1p::RFP] III; dsl-3(ok3411) IV* | This study |
| IS1894 | *wyIs75[unc-47p::DsRed; exp-1p::GFP; odr-1p::RFP] III; dsl-3(ok3411) IV; sel-12(ar171) unc-1(e538) X* | This study |
| IS1890 | *syIs63[cog-1::gfp;unc-119(+)] IV; dsl-4(ok1020)X* | This study |
| IS1958 | *rrf-3(pk1426) II; syIs63[cog-1::gfp;unc-119(+)] IV; dsl-4(ok1020)X* | This study |
| IS1872 | *wyIs75[unc-47p::DsRed; exp-1p::GFP; odr-1p::RFP] III; dsl-5(ok588) IV* | This study |
| IS1873 | *wyIs75[unc-47p::DsRed; exp-1p::GFP; odr-1p::RFP] III; dsl-5(ok588) IV; sel-12(ar171) unc-1(e538) X* | This study |
| IS1874 | *wyIs75[unc-47p::DsRed; exp-1p::GFP; odr-1p::RFP] III; dsl-6(ok2265) IV* | This study |
| IS1875 | *wyIs75[unc-47p::DsRed; exp-1p::GFP; odr-1p::RFP] III; dsl-6(ok2265) IV; sel-12(ar171) unc-1(e538) X* | This study |
| IS1887 | *osm-7(n1515) III; syIs63[cog-1::gfp;unc-119(+)] IV* | This study |
| IS1888 | *osm-7(n1515) III; syIs63[cog-1::gfp;unc-119(+)] IV; sel-12(ar171) unc-1(e538) X* | This study |
| IS1924 | *syIs63[cog-1::gfp;unc-119(+)] IV; osm-11(rt192) X* | This study |
| IS1960 | *rrf-3(pk1426) II; syIs63[cog-1::gfp;unc-119(+)] IV; osm-11(rt192)X* | This study |
| IS1876 | *dos-1(ok2398) III; syIs63[cog-1::gfp;unc-119(+)] IV* | This study |

| | | |
|---|---|---|
| IS1877 | *dos-1(ok2398) III; syIs63[cog-1::gfp;unc-119(+)] IV; sel-12(ar171) unc-1(e538) X* | This study |
| IS1889 | *dos-2(tm4515) II; syIs63[cog-1::gfp;unc-119(+)] IV* | This study |
| IS1878 | *dos-2(tm4515) II; syIs63[cog-1::gfp;unc-119(+)] IV; sel-12(ar171) unc-1(e538) X* | This study |
| IS1879 | *dos-3(tm4899) III; syIs63[cog-1::gfp;unc-119(+)] IV* | This study |
| IS1881 | *dos-3(tm4899) III; syIs63[cog-1::gfp;unc-119(+)] IV; sel-12(ar171) unc-1(e538) X* | This study |
| Transgenic strains used to assess ligand availability over time | | |
| IS2164 | *syIs63[cog-1::gfp;unc-119(+)] IV; fpEx713[col-34p::lin-12cDNA(n941); myo-2p::mCherry] ** | This study |
| IS2166 | *syIs63[cog-1::gfp;unc-119(+)] IV; fpEx714[col-34p::lin-12cDNA(n941); myo-2p::mCherry] ** | This study |
| IS2169 | *syIs63[cog-1::gfp;unc-119(+)] IV; fpEx717[col-34p::lin-12cDNA(n941); myo-2p::mCherry] ** | This study |
| IS2193 | *syIs63[cog-1::gfp;unc-119(+)] IV; fpEx729[col-34p::lin-12cDNA(ΔANK); myo-2p::mCherry] ** | This study |
| IS2195 | *syIs63[cog-1::gfp;unc-119(+)] IV; fpEx731[col-34p::lin-12cDNA(ΔANK); myo-2p::mCherry] ** | This study |
| IS2196 | *syIs63[cog-1::gfp;unc-119(+)] IV; fpEx732[col-34p::lin-12cDNA(ΔANK); myo-2p::mCherry] ** | This study |
| IS2161 | *syIs63[cog-1::gfp;unc-119(+)] IV; fpEx710[col-34p::lin-12cDNA(n137); myo-2p::mCherry] ** | This study |
| IS2162 | *syIs63[cog-1::gfp;unc-119(+)] IV; fpEx711[col-34p::lin-12cDNA(n137); myo-2p::mCherry] ** | This study |
| IS2163 | *syIs63[cog-1::gfp;unc-119(+)] IV; fpEx712[col-34p::lin-12cDNA(n137); myo-2p::mCherry] ** | This study |
| IS2130 | *syIs63[cog-1::gfp;unc-119(+)] IV; fpEx689[col-34p::lin-12cDNA; myo-2p::mCherry] ** | This study |
| IS2132 | *syIs63[cog-1::gfp;unc-119(+)] IV; fpEx691[col-34p::lin-12cDNA; myo-2p::mCherry] ** | This study |
| IS2133 | *syIs63[cog-1::gfp;unc-119(+)] IV; fpEx692[col-34p::lin-12cDNA; myo-2p::mCherry] ** | This study |
| Expression of NODE-like complex members in $Y^{prog}$ and $DA9^{Prog}$ | | |
| MH1337 | *kuIs34[sem-4::gfp; unc-119(+)]; unc-119(ed3)* | Grant *et al.*, 2000 |
| IS2213 | *fpEx476[GFP::ceh-6 fosmid; odr-1::RFP] *; ceh-6(gk665)* | This Study |
| IS2239 | *fpEx756[GFP::sox-2 fosmid; odr-1::rfp]* ; sox-2(ot640) X* | This Study |

| | | |
|---|---|---|
| IS2857 | *fpEx974 [NLS :GFP ::T2A ::SEM-4 Fosmid; dsred::coelomocyte]*; fpIs67[itr-1::mCherry; odr-1::GFP]*; syIs63[cog-1::gfp;unc-119(+)] IV* | This Study |
| IS3 | *bxIs7[egl-5::gfp; lin-15(+)]* | Jarriault *et al.*, 2008 |
| IS2540 | *fpIs88[hlh-16::GFP; pRF4]** | This Study |
| IS2807 | *fpEx876[lin-12p::ICL::lin-12UTR (0,5ng); myo-2::mCherry (2ng)]* ; fpEx974[NLS::GFP::T2A::sem-4 Fosmid (50ng) ; DsRed::Coelomocyte (50ng)] ** | This Study |
| IS2802 | *fpEx876[lin-12p::ICL::lin-12UTR (0,5ng); myo-2::mCherry (2ng)]* ; fpIs100(egl-5(1,3kb)delta pes10::mkate::unc-54 3'UTR; cc:GFP from fpEx970]* ; fpIs88 [hlh-16::GFP translational; pRF4]** | This Study |
| Extra PDA in *lin-12(gf)* come from a cell that first adopts Y fate | | |
| IS2528 | *lin-12(n950) III ; kuIs34[sem-4::gfp; unc-119(+)]* | This Study |
| Regulation of *lin-12* expression is at the transcriptional level | | |
| IS2443 | *fpEx875[lin-12p::NICDGFP::lin-12UTR; myo-2::mCherry] *; syIs63[cog-1::gfp;unc-119(+)] IV ; fpIs67[odr-1:GFP; itr-1p::mcherry]* | This Study |
| IS2444 | *fpEx876[lin-12p::NICDGFP::lin-12UTR; myo-2::mCherry] *; syIs63[cog-1::gfp;unc-119(+)] IV ; fpIs67[odr-1:GFP; itr-1p::mcherry]* | This Study |
| IS2445 | *fpEx877[lin-12p::NICDGFP::lin-12UTR; myo-2::mCherry] *; syIs63[cog-1::gfp;unc-119(+)] IV ; fpIs67[odr-1:GFP; itr-1p::mcherry]* | This Study |
| IS2446 | *fpEx878[lin-12p::NICDGFP::lin-12UTR; myo-2::mCherry] *; syIs63[cog-1::gfp;unc-119(+)] IV ; fpIs67[odr-1:GFP; itr-1p::mcherry]* | This Study |
| IS2447 | *fpEx879[lin-12p::NICDGFP::lin-12UTR; myo-2::mCherry] *; syIs63[cog-1::gfp;unc-119(+)] IV ; fpIs67[odr-1:GFP; itr-1p::mcherry]* | This Study |
| IS2504 | *fpEx891[lin-12p(ΔR1)::NICDGFP::lin-12UTR; myo-2::mCherry] *; syIs63[cog-1::gfp;unc-119(+)] IV ; fpIs67[odr-1:GFP; itr-1p::mcherry]* | This Study |
| IS2505 | *fpEx892[lin-12p(ΔR1)::NICDGFP::lin-12UTR; myo-2::mCherry] *; syIs63[cog-1::gfp;unc-119(+)] IV ; fpIs67[odr-1:GFP; itr-1p::mcherry]* | This Study |
| IS2506 | *fpEx893[lin-12p(ΔR1)::NICDGFP::lin-12UTR; myo-2::mCherry] *; syIs63[cog-1::gfp;unc-119(+)] IV ; fpIs67[odr-1:GFP; itr-1p::mcherry]* | This Study |
| IS2507 | *fpEx894[lin-12p(ΔR1)::NICDGFP::lin-12UTR; myo-2::mCherry] *; syIs63[cog-1::gfp;unc-119(+)] IV ; fpIs67[odr-1:GFP; itr-1p::mcherry]* | This Study |
| IS2508 | *fpEx895[lin-12p(ΔR1)::NICDGFP::lin-12UTR; myo-2::mCherry] *; syIs63[cog-1::gfp;unc-119(+)] IV ; fpIs67[odr-1:GFP; itr-1p::mcherry]* | This Study |
| IS2509 | *fpEx896[lin-12p(ΔR2)::NICDGFP::lin-12UTR; myo-2::mCherry] *; syIs63[cog-1::gfp;unc-119(+)] IV ; fpIs67[odr-1:GFP; itr-1p::mcherry]* | This Study |
| IS2510 | *fpEx897[lin-12p(ΔR2)::NICDGFP::lin-12UTR; myo-2::mCherry] *; syIs63[cog-1::gfp;unc-119(+)] IV ; fpIs67[odr-1:GFP; itr-1p::mcherry]* | This Study |

| | | |
|---|---|---|
| IS2511 | *fpEx898[lin-12p(ΔR2)::NICDGFP::lin-12UTR; myo-2::mCherry] *; syIs63[cog-1::gfp;unc-119(+)] IV ; fpIs67[odr-1:GFP; itr-1p::mcherry]* | This Study |
| IS2512 | *fpEx899[lin-12p(ΔR2)::NICDGFP::lin-12UTR; myo-2::mCherry] *; syIs63[cog-1::gfp;unc-119(+)] IV ; fpIs67[odr-1:GFP; itr-1p::mcherry]* | This Study |
| IS2513 | *fpEx900[lin-12p(ΔR2)::NICDGFP::lin-12UTR; myo-2::mCherry] *; syIs63[cog-1::gfp;unc-119(+)] IV ; fpIs67[odr-1:GFP; itr-1p::mcherry]* | This Study |
| Strains to assess Notch level effect on Y-to-PDA timing | | |
| IS4210 | *fpEx686[mig-13p::mCherry (8ng); myo-2::Mcherry (2ng); pBS (190ng)] * ; syIs63[cog-1::gfp;unc-119(+)] IV* | This Study |
| IS4211 | *fpEx686[mig-13p::mCherry (8ng); myo-2::Mcherry (2ng); pBS (190ng)] *; unc-32(e189) III lin-12(n676n930) III ; syIs63[cog-1::gfp;unc-119(+)] IV* | This Study |
| IS3 | *bxIs7[egl-5::gfp; lin-15(+)]* | Jarriault *et al.*, 2008 |
| IS764 | *unc-32(e189) III lin-12(n676n930) III ; bxIs7[egl-5::gfp; lin-15(+)] I* | This Study |
| IS4239 | *unc-32(e189) III lin-12(n676n930) III ; hlh-16(syb683[GFP::linker::hlh-16]) I* ; fpEx686[mig-13p::mCherry (8ng); myo-2::Mcherry (2ng); pBS (190ng)] ** | This Study |
| IS4257 | *hlh-16(syb683[GFP::linker::hlh-16]) I *; fpEx686[mig-13p::mCherry (8ng); myo-2::Mcherry (2ng); pBS (190ng)] ** | This Study |
| IS4252 | *fpEx686[mig-13p::mCherry (8ng); myo-2::Mcherry (2ng); pBS (190ng)] *; ujIs113 [pie-1p::mCherry::H2B::pie-1 3'UTR + nhr-2p::mCherry::his-24::let-858 3'UTR + unc-119(+)] II ; ngn-1(dev137([mNeonGreen::ngn-1]) IV ; unc-32(e189) III lin-12(n676n930) III* | This Study |
| IS4262 | *fpEx686[mig-13p::mCherry (8ng); myo-2::Mcherry (2ng); pBS (190ng)] *; ujIs113 [pie-1p::mCherry::H2B::pie-1 3'UTR + nhr-2p::mCherry::his-24::let-858 3'UTR + unc-119(+)] II ; ngn-1(dev137([mNeonGreen::ngn-1]) IV* | This Study |
| IS4247 | *unc-32(e189) III lin-12(n676n930) III ; muIs62[Pmig-13-mig-13::GFP + lin-15(+)] ; nsIs913[ngn-1p::myr-mKate2 (5 ng/ul); myo-2p::mCherry (2.5 ng/ul); pBlueScript (92.5 ng/ul)] IV (might contain mig-13(mu225) X lin-15(n765ts) X)* | This Study |
| IS4248 | *muIs62[Pmig-13-mig-13::GFP + lin-15(+)] ; nsIs913[ngn-1p::myr-mKate2 (5 ng/ul); myo-2p::mCherry (2.5 ng/ul); pBlueScript (92.5 ng/ul)] IV (might contain mig-13(mu225) X lin-15(n765ts) X)* | This Study |
| IS563 | *lin-12(n302) III ; syIs63[cog-1::gfp;unc-119(+)] IV* | This Study |
| IS2118 | *lin-12(n950) III ; fpEx682[mig-13p::mCherry (8ng); myo-2::Mcherry (2ng); pBS (190ng)]* ; syIs63[cog-1::gfp;unc-119(+)] IV* | This Study |
| IS2444 | *fpEx876[lin-12p::ICL::lin-12UTR (0,5ng); myo-2::mCherry (2ng)] * ; syIs63[cog-1::gfp;unc-119(+)] IV ; fpIs67[odr-1:GFP; itr-1p::mcherry]** | This Study |
| IS3873 | *hlh-16(syb683[GFP::linker::hlh-16]) I* ; fpIs130[pSJ1007-10µg/ml; pBSK+-200ng/ml]N2 ; fpEx876[lin-12p::ICL::lin-12UTR (0,5ng); myo-2::mCherry (2ng)]** | This Study |

### SI Table 3: Extrachromosomal arrays generated for this study

| Transgene | Co-injection marker | Plasmid | Injection in |
|---|---|---|---|
| *fpEx683, fpEx684, fpEx686* | myo-2::mCherry (2ng/µL) | pCM327 – mig-13p::mCherry::unc-10 3' UTR (8 ng/µL) (a gift from the Shen laboratory) | *syIs63[cog-1::gfp;unc-119(+)] IV* |
| *fpEx682* | myo-2::mCherry (2ng/µL) | pCM327 – mig-13p::mCherry::unc-10 3' UTR (8 ng/µL) (a gift from the Shen laboratory) | *lin-12(n950) III; syIs63[cog-1::gfp;unc-119(+)] IV* |
| *fpEx664, fpEx665, fpEx666* | myo-2::mCherry (2ng/µL) | pSJ3171 - hsp-16.2::NICDGFP::unc-54 3′UTR (10ng/µL) | *hsf-1(sy441)I* |
| *fpEx84, fpEx85, fpEx86, fpEx87, fpEx88* | myo-2p::GFP (5ng/µL) | pSJ6003 - egl-5(6,2kb)Δpes10p::NICDGFP::SL2::mCherry::unc-54 3′UTR (20ng/µL) | *syIs63[cog-1::gfp;unc-119(+)] IV* |
| *fpEx217, fpEx218, fpEx219, fpEx220, fpEX221* | myo-2p::GFP (2ng/µL) | pSJ3173 – col-34p::NICDGFP::unc-54 3′UTR (20ng/µL) | *syIs63[cog-1::gfp;unc-119(+)] IV.* |
| *fpEx334, fpEx335, fpEx336, fpEx337, fpEx338* | myo-2p::GFP (2ng/µL) | pSJ3169 – lin-48p::NICDGFP::SL2::mCherry::unc-54 3′UTR (20ng/µL) | *syIs63[cog-1::gfp;unc-119(+)] IV.* |
| *fpEx459, fpEx460, fpEx461, fpEx462, fpEx463* | myo-2p::mCherry (2ng/µL) | pSJ3162 – egl-20p::NICDGFP::SL2::mCherry::unc-54 3′UTR (20ng/µL) | *syIs63[cog-1::gfp;unc-119(+)] IV* |
| *fpEx713, fpEx714, fpEx717* | myo-2p::mCherry (2ng/µL) | pSJ3222 - col-34p::lin-12cDNA(n941)::unc-54 3′UTR (10ng/µL) | *syIs63[cog-1::gfp;unc-119(+)] IV* |
| *fpEx729, fpEx73, fpEx732* | myo-2p::mCherry (2ng/µL) | pSJ3223 - col-34p::lin-12cDNA(ΔANK)::unc-54 3′UTR (10ng/µL) | *syIs63[cog-1::gfp;unc-119(+)] IV* |
| *fpEx710, fpEx711, fpEx712* | myo-2p::mCherry (2ng/µL) | pSJ3218 - col-34p::lin-12cDNA(n137)::unc-54 3′UTR (10ng/µL) | *syIs63[cog-1::gfp;unc-119(+)] IV* |
| *fpEx689, fpEx691, fpEx692* | col-34p::lin-myo-2p::mCherry (2ng/µL) | pSJ3215 - col-34p::lin-12cDNA::unc-54 3′UTR (10ng/µL) | *syIs63[cog-1::gfp;unc-119(+)] IV* |
| *fpEx875, fpEx876, fpEx877, fpEx878, fpEx879* | myo-2p::mCherry (2ng/µL) | pSJ3212 – lin-12p::NICDGFP::lin-12UTR (0.5ng/µL) | *syIs63[cog-1::gfp;unc-119(+)] IV; fpIs67[itr-1p::mCherry; odr-1p::GFP]* |
| *fpEx891, fpEx892, fpEx893, fpEx894, fpEx895* | myo-2p::mCherry (2ng/µL) | pSJ3240 – lin-12p(ΔR1)::NICDGFP::lin-12UTR (0.5ng/µL) | *syIs63[cog-1::gfp;unc-119(+)] IV; fpIs67[itr-1p::mCherry; odr-1p::GFP]* |
| *fpEx896, fpEx897, fpEx898, fpEx899, fpEx900* | myo-2p::mCherry (2ng/µL) | pSJ3242 – lin-12p(ΔR2)::NICDGFP::lin-12UTR (0.5ng/µL) | *syIs63[cog-1::gfp;unc-119(+)] IV; fpIs67[itr-1p::mCherry; odr-1p::GFP]* |

| | | | |
|---|---|---|---|
| *fpEx476* | myo-2p::mCherry (2ng/µL) | Fosmid 9347172996193398_D03 = ceh-6::GFP::3xFLAG (20ng/µL)<br>(a gift from Mihail Sarov) | *ceh-6(ok3388) I / hT2[qIs48] strain* |
| *fpEx756* | odr-1::rfp (50ng/µL) | Fosmid WRM0626aE02 = GFP::sox-2 (10ng/µL) | *sox-2(ot640) X* |
| *fpEx828* | myo-2::GFP (3ng/µL) | pSJ821 - hlh-16p::mcherry::hlh-16::hlh-16 3'UTR (20ng/µL) | *hlh16(fp12 ) I ; syIs63[cog-1::gfp;unc-119(+)] IV* |
| *fpEx830* | myo-2::GFP (3ng) | pSJ823 - col-34p::mcherry::hlh-16::hlh-16 3'UTR (20ng/µL) | *hlh16(fp12 ) I ; syIs63[cog-1::gfp;unc-119(+)] IV* |
| *fpEx929* | myo-2::GFP (2ng/µl) | pSJ6334 – 5.6kb ceh-6p::GFP::3.6kb ceh-6 3'UTR (10ng/µl) | N2 |
| *fpEx974* | dsred::coelomocyte (50ng/µL) | Fosmid WRM0638dE10, modified as NLS::GFP ::T2A::SEM-4 (50ng/µL) | N2 |

### SI Table 4: Integrated arrays generated for this study

| Integrated array | Extrachromosomal array | Plasmid name & gene | genetic background |
|---|---|---|---|
| *fpIs130* | *fpEx1251* | pSJ1007 = egl-5p(6 kb)::2xNLS::mCherry::unc-54 3'utr (10ng/µL) | N2 |
| *fpIs88* | *otEx4503* from Bertrand et al. 2011 | [hlh-16::GFP; pRF4] | N2 |
| *fpIs100* | *fpEx970* | pSJ834 = egl-5(1,3kb)Δpes10::mkate::unc-54 3'UTR(20ng/µL); cc::GFP (50ng/µL) | N2 |
| *fpIs10; fpIs11* | *fpEx30* | pSJ503 = exp-1p::mCherry(10ng/µL); myo-2p::GFP (10ng/µL) | N2 |
| *fpIs51, fpIs54* | *fpEx497* | pSJ6003 = egl-5(6,2kb)Δpes10p::NICD::SL2::mCherry (20ng/µL) | N2 |
| *fpIs67* | *wyEx1902* from Teichmann & Shen, 2011 | [itr-1pB::mCherry; odr-1p::GFP] | N2 |

### SI Table 5: Oligonucleotides used

| Oligo name | Sequence | Use |
|---|---|---|
| BDM314 | AACGGTACCAGAAAAAATGGTTGTTCTGATGTTAGGAGCATTACC | pSJ3177 and pSJ6003 cloning |
| BDM315 | TTGGTACCTCAAAAATAATGAGCTGGTTCGGAGTATCG | pSJ3177 and pSJ6003 cloning |
| BDN519 | ACAGCATGCGACATGTAAAGTACATCCGTTACATC | pSJ3173 cloning |
| BDN521 | CCCCCCGGGTGTATGCAGTGGTGGTTTGG | pSJ3173 cloning |
| BDT599 | ACATGCATGCGGATCCAAAAAACCTGCATTTTTTTCAG | pSJ3169 cloning (*lin-48* promoter) |
| BDT600 | CCCCCCGGGCTGAAATTGAGCAGAGCTGAAAATTTTTG | pSJ3169 cloning (*lin-48* promoter) |
| Ceh-6pF | ATAAGAATgcggccgcCGTGTTGCTTTAGCACTTCTCCATCCCTTC | pSJ6334 (*ceh-6* |

| | | |
|---|---|---|
| | | transcriptional reporter) |
| Ceh-6pR | ATAGTTTAgcggccgcCAGTTGGGAAGTCCAGGAGCAACGGGGTG | pSJ6334 (*ceh-6* transcriptional reporter) |
| Ceh-6UTRf | TTTTTTGTGATGCGTATTGATGTAGC | pSJ6334 (*ceh-6* transcriptional reporter) |
| Ceh-6UTRr | GTCGACACAGAAACTACGCAAAATC | pSJ6334 (*ceh-6* transcriptional reporter) |
| mcm239F | TAGAGGATCCccgGGGATTGGCC | pSJ834 cloning (mkate insertion) |
| mcm239R | GATATTATACATATTTCATAAAGCCAACC | pSJ834 cloning (mkate insertion) |
| oCG521 | CGAGCTCAGAAAAAATGACAGCACCGAAAAAAAAGCGAAAAGTTCCA GCTGAGAAGATGACCGCTCCAAAGAAGAAACGCAAAGTA | pSJ1007 cloning |
| oCG522 | CCGGTACTTTGCGTTTCTTCTTTGGAGCGGTCATCTTCTCAGCTGGAACT TTTCGCTTTTTTTTCGGTGCTGTCATTTTTTCTGAGCTCGGTAC | pSJ1007 cloning |
| TD004 | GCGGCCGCGCTGTCTCATCCTACTTTCACC | pSJ3169 cloning (*SL2::mCherry*) |
| TD005 | GCGGCCGCCTACTTATACAATTCATCCATGCC | pSJ3169 cloning (*SL2::mCherry*) |
| TD061 | AAAGCATGCGAAGTCATCCTACTAACTAACAATATGACGC | pSJ3162 cloning (*egl-20* promoter) |
| TD062 | AAACCCGGGTATTTCTGAAATTGAGATGTTTTAGAATTTC | pSJ3162 cloning (*egl-20* promoter) |
| TD065 | GACTCAACTCATCTGACACCTCC | pSJ3103 cloning |
| TD066 | GGGTCGAGTTACTTTTCTTGAAGG | pSJ3103 cloning |
| TD088 | taatacgactcactatagggATGCCTTCCACAAGGAGACAAC | *sel-12* gDNA PCR amplification for RNAi experiments |
| TD089 | taatacgactcactatagggGAGATCGCTCAAGATATAATCGAAAAG | *sel-12* gDNA PCR amplification for RNAi experiments |
| TD094 | AAAAGGTACCATGCGGATCCCTACGATTTG | pSJ3215 cloning |
| TD095 | TTTTGGTACCTCAAAAATAATGAGCTGGTTCGG | pSJ3215 cloning |
| TD101 | GTGTTGTTGACTCAATATTTGCAAGGCTTGC | pSJ3218 cloning |

| | | |
|---|---|---|
| TD102 | GCAAGCCTTGCAAATATTGAGTCAACAACAC | pSJ3218 cloning |
| TD105 | GGATTCGGTGGGAAATAGTGTGACGAGCCATTG | pSJ3222 cloning |
| TD106 | CAATGGCTCGTCACACTATTTCCCACCGAATCC | pSJ3222 cloning |
| TD113 | CCAGAACGAGAATATTCAATGGATC | pSJ3223 cloning |
| TD114 | AGGTTCAGGTTCAGTTGGAATTTG | pSJ3223 cloning |
| TD096 | CTCAACAGACTTTGCTCAATTTCAAAAAATGGTTGTTCTGATGTTAG GAGCATTAC | pSJ3212 cloning |
| TD097 | GGAATTTAAATAATAAATGACGATTGTTCAGAAGATGTACCGAGCT CGGATCCACTAGTAAC | pSJ3212 cloning |
| TD136 | ACAGTAACAGACACCTGTGCTCC | pSJ3240 cloning |
| TD137 | TATTGTTAATAAATGAGTGTAACATTTAAG | pSJ3240 cloning |
| TD138 | ATTAATGATAATGCAAAAGCTACCAGG | pSJ3242 cloning |
| TD139 | TGTGTCAGTTTTAGAGTTTTATTTCTG | pSJ3242 cloning |
